# AbBFN2: A flexible antibody foundation model based on Bayesian Flow Networks

**DOI:** 10.1101/2025.04.29.651170

**Authors:** Bora Guloglu, Miguel Bragança, Alex Graves, Scott Cameron, Timothy Atkinson, Liviu Copoiu, Alexandre Laterre, Thomas D. Barrett

## Abstract

Antibody engineering is marked by diverse data and desiderata, making it a prime candidate for multi-objective design, but is commonly tackled as a series of sequential optimisation tasks. Here, we present AbBFN2, a generative antibody foundation model trained on paired antibody sequences as well as genetic and biophysical metadata. This is achieved using the Bayesian Flow Network paradigm, which allows unified modelling of diverse data sources and flexible conditional generation at inference time. By virtue of its rich set of features and architectural flexibility, AbBFN2 can be adapted to a number of tasks commonly tackled by individual models, consolidating traditional computational pipelines into a single step. We demonstrate the adaptability of AbBFN2 using sequence inpainting, humanisation, biophysical property optimisation, and conditional *de novo* library generation of antibodies with rare attributes as example tasks. By removing the need for task-specific training, we hope that AbBFN2 will accelerate machine learning-based antibody design and development workflows.

## Introduction

Antibodies constitute an essential component of the adaptive immune response. These proteins demonstrate remarkable specificity and affinity for their antigens, binding to them and recruiting the downstream immune response [1]. Due to their ability to recognise virtually any foreign target, antibodies have also shown remarkable success as a therapeutic modality, with a market size of $252.6 billion in 2024, which is further projected to increase to $0.5 trillion by 2029 [2]. Indeed, therapeutic antibodies make up a major share of new clinical trials, with at least 12 antibodies entering either the US or EU market every year since 2020 [3].

Not all antibodies make good therapeutic candidates. A tractable sequence must not only bind its antigen without off-target effects, but it must also be free of developability issues such as aggregation propensity, low expression levels, or poor solubility [4, 5]. As such, a discovery campaign usually starts with an initial binder, which is then optimised in an iterative process to remove any potential liabilities whilst maintaining or improving the affinity of the antibody for its intended target [6]. A plethora of computational and experimental approaches are used to form a development pipeline to achieve this [7, 8]. Since many of the methods are agnostic to properties optimised by upstream steps, such pipelines run the risk of deteriorating one property while optimising another. Similarly, the design of antibody libraries from which initial candidates can later be isolated is often a laborious process. This usually involves a number of filtering steps in order to decrease the occurrence of unfavourable properties in the library [9], or to enrich the library in attributes that are known to be important for interactions with the target antigen [10, 11]. The antibody design problem is further complicated by the vast search space that one must navigate, with the theoretical diversity of naive antibody sequences estimated to be in the order of 10^10^ *−* 10^11^ [12, 13]. Indeed, a number of forward-looking studies have conceptualised antibody design as a multi-objective optimisation task, where all relevant properties should be optimised in a concerted manner [14–16].

Here, we present AbBFN2, a flexible tool for *de novo* sequence generation, analysis, and antibody design based on the Bayesian Flow Network (BFN) [17, 18] paradigm. BFNs model parameters of distributions over data rather than the data itself, and are able to work with discrete and continuous data types, providing a unifying framework with which to model these distinct data modalities. By incorporating information describing the species, genetic lineages, and therapeutically relevant properties pertaining to an antibody alongside its sequence, AbBFN2 allows for concurrent optimisation of a number of relevant properties. AbBFN2 jointly models antibody sequences and information describing the species, genetic lineages, and therapeutically relevant properties, providing a rich grammar with which to guide generation. Concretely, by conditionally generating subsets of variables given user defined context over the others, AbBFN2 is able to realise an array of tasks at test time, without further training. Here, we demonstrate this flexibility using a number of diverse tasks normally tackled by separate models: sequence labelling, sequence humanisation and redesign, and conditional *de novo* library generation.

## Results

AbBFN2 is a 800M parameter model trained on the known paired antibody sequences in the OAS database [19, 20] (Fig. 1). This dataset covers approximately 2M human, mouse, and rat antibody sequences, with a heavy bias in favour of sequences coming from human repertoires (95 % human). Additionally, the training process involved annotation of the sequences with the species, CDR length, and genetic labels (derived from the nucleotide sequences, rather than on the amino acid level), as well as surface properties derived from the Therapeutic Antibody Profiler (TAP), [5, 23] (Fig. 1a). During sampling, the model concurrently generates labels for all of these attributes alongside the sequence (Fig. 1b). Details regarding the training and sampling procedures are provided in the Methods section.

**Figure 1.**
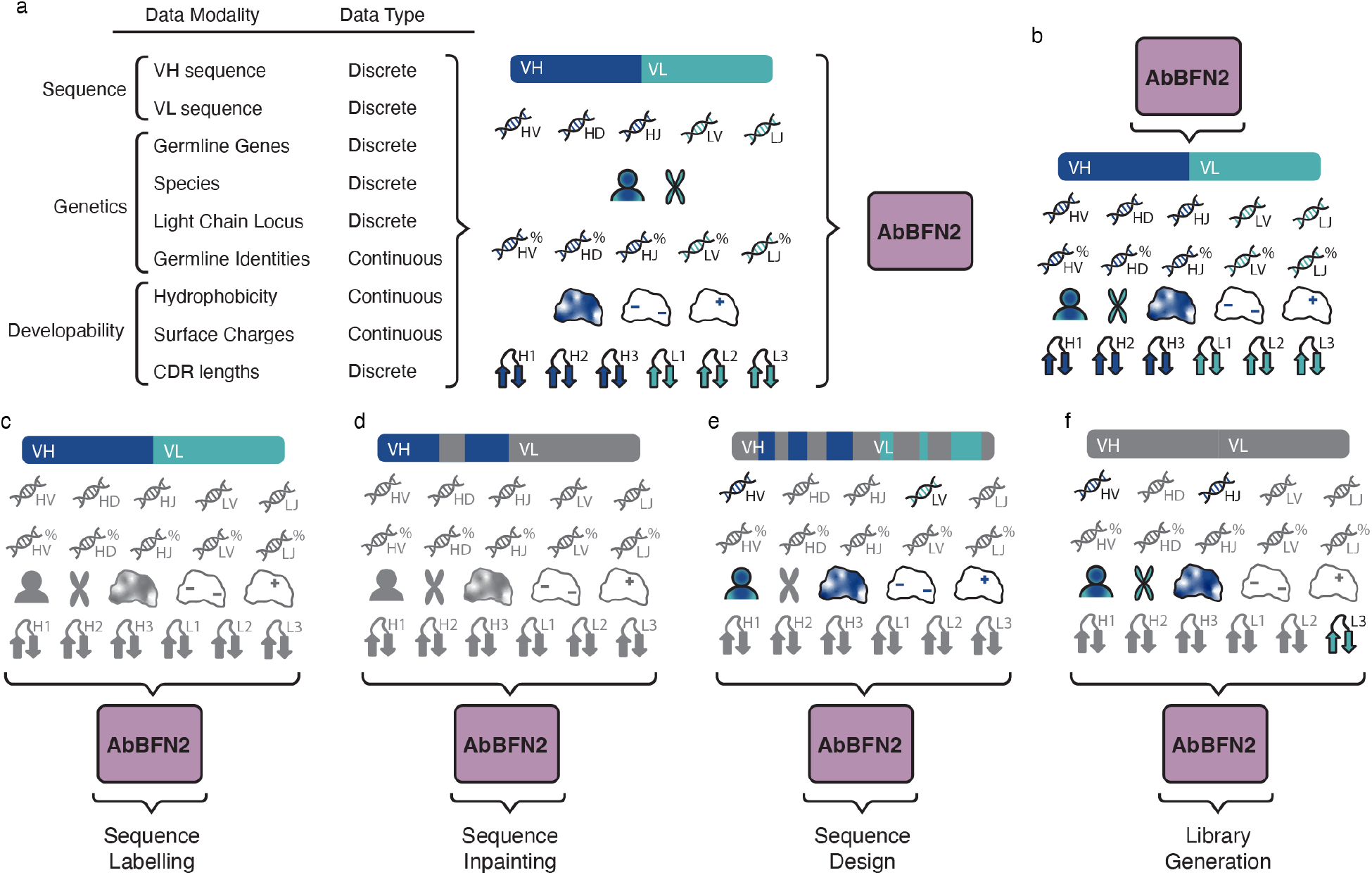
Overview of AbBFN2. **(a)** shows the data that the model is trained on. Namely, we extracted all paired antibody chains from OAS [19, 20] alongside their genetic annotations derived from IgBLASTn [21], structurally modelled them using ImmuneBuilder [22], and derived therapeutic properties from these models. We then included all metadata (genetic lineages, developability scores, etc.) as well as the full sequence as data modalities to train AbBFN2. The data modes of AbBFN2 provide the user with a comprehensive representation of the properties of antibodies which can be used to guide the generative process. **(b)** During inference, AbBFN2 generates all data modes concurrently. When no conditioning information is provided, this is equivalent to unconstrained *de novo* generation. **(c-f)** show different use-cases of AbBFN2 based on arbitrary combinations of conditioning information, allowing the model to handle a variety of tasks. As examples, we show sequence labelling, sequence inpainting, sequence design, and library generation, respectively.

### AbBFN2 successfully learns relationships across antibody sequence, genetics, and biophysical properties

The core aim of AbBFN2 is to provide a model capable of highly tunable conditional generation. To this end, we included 31 additional data modes spanning genetic and biophysical attributes of each training example in addition to the 14 data modes encoding the antibody sequence (Fig. 1a). Having learned the underlying joint distribution, AbBFN2 concurrently generates all 45 data modes at inference time. A prerequisite for effective generation by such multi-modal models is faithful capturing of the underlying marginal distributions. In order to assess the extent to which AbBFN2 has learned the relationships between the modelled attributes of of natural, paired antibodies, we unconditionally generated 10 000 samples (Fig. 1b), as described in the Methods section and examined the marginal and conditional distributions of the different data modes, comparing them to a hold-out set of the same size. In this section, we show that AbBFN2 has learned the joint distribution over the modelled data modes by assessing a number of relevant dependencies.

### Analysis of generated CDR loops

As the CDR loops are the most functionally relevant and variable regions of an antibody, we first analyse the sequence-based and structural properties of generated CDR loops, comparing them to expected natural distributions and known variability. We find that generated CDR sequences match the natural population in terms of length (Fig. 2a-f) and amino acid propensities, both on a per-position basis (Fig. 2g, Fig. S1, Fig. S2, Fig. S3, Fig. S4) and averaged across each CDR loop (Fig. S5). Where we do observe small deviations in amino acid usage (< 1.5 %), these substitutions are often biophysically justified as they are expected to preserve overall function; for example the enrichment of Lys over Arg residues in V_H_ IMGT position 106, both of which are positively charged (Fig. 2g). We describe full details and extended analysis in F.1.

**Figure 2.**
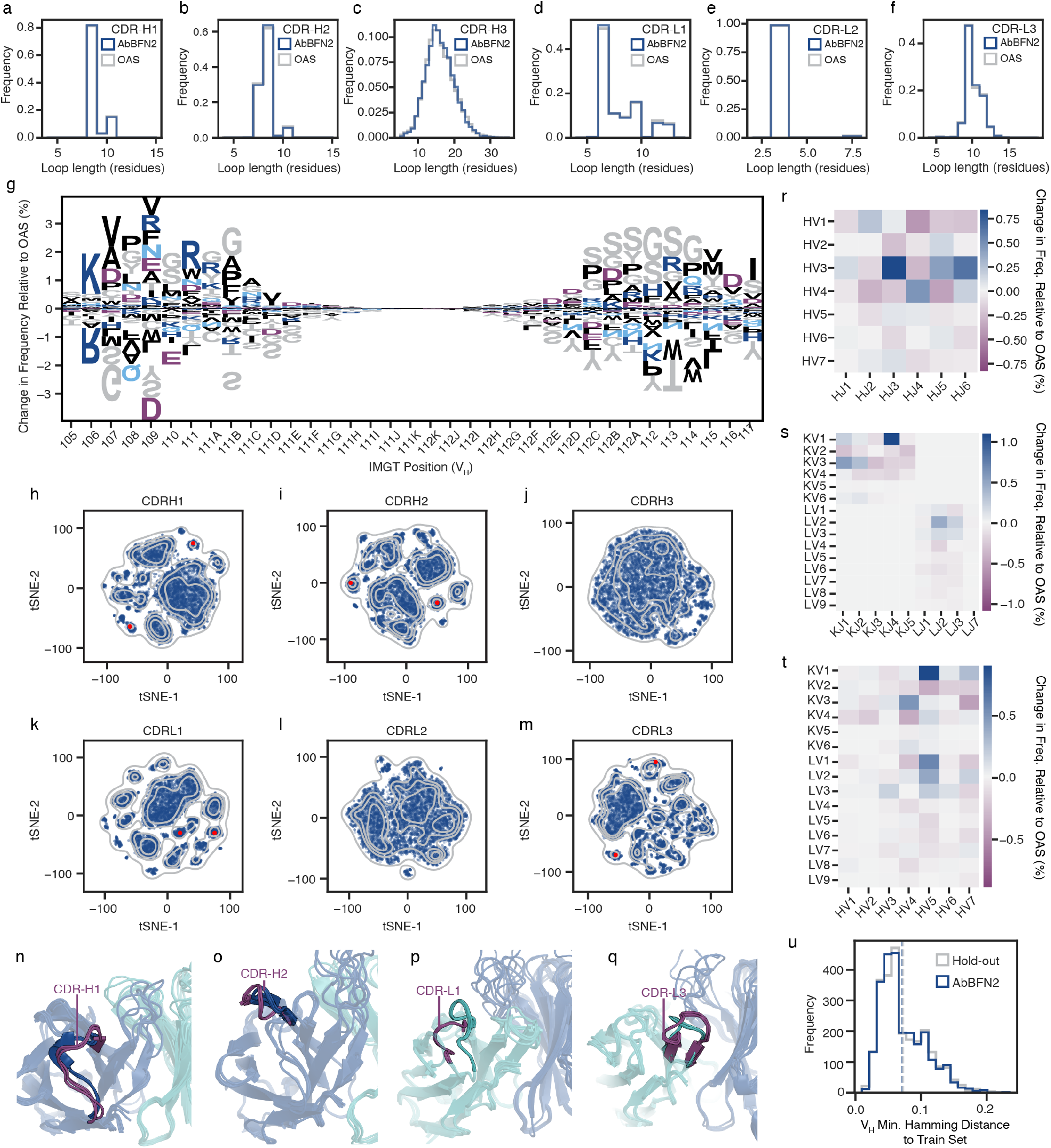
Unconditional generation of antibody sequences using AbBFN2. **(a-f)** show distributions of CDR loop lengths from held-out (grey) and generated (navy) samples. CDRs are defined according to the IMGT scheme. **(g)** shows position-specific deviations of amino acid frequencies of generated sequences when compared to the held out data across CDR-H3. Residues are coloured according to their physicochemical properties. Positive values correspond to amino acids that are overrepresented in generated samples, and *vice versa*. **(h-m)** shows structural comparisons of generated and held out samples, folded using ABodyBuilder2. Briefly, for each CDR, samples were pooled, aligned on the CDR anchors and pairwise loop structural distances were calculated using dynamic time warping. Pairwise distances were then used to create two-dimensional tSNE embeddings. Grey contour plots refer to kernel density estimates of held-out data; navy scatter plots refer to generated samples. Red points correspond to samples depicted in **(n-q)**, with one cluster depicted in navy/cyan, and another depicted in purple for each CDR. **(r-t)** refer to deviations in germline gene pairing propensities of generated samples compared to held out data. Positive values correspond to pairings that are overrepresented in generated samples, and *vice versa*. **(u)** shows distributions of Hamming distances between the held-out data (grey) and generated samples (blue) and a random sample of 10,000 sequences from the training set. Briefly, each sequence was aligned and padded to the most commonly found 200 IMGT positions, before distance calculation and normalisation. For each generated or held-out sequence, the smallest Hamming distance to the train set is reported. Dashed lines correspond to sample means.

### CDR structures

In order to examine the CDR loops as a whole and in the context of the FW regions surrounding them, we structurally modelled all 10 000 generated sequences and 10 000 held-out sequences using ImmuneBuilder [22] (9995 and 10 000 were successfully modelled, respectively) and clustered the CDR loop structures using dynamic time warping (DTW) as described in the Methods section to perform pairwise comparisons (Fig. 2h-q). Notably, we were able to recover the same natural structural clusters in both datasets for the CDR-H1/H2/L1/L3 loops in a manner similar to previous studies (Fig. 2h, Fig. 2i, Fig. 2k, Fig. 2m, respectively) [24–26]. The CDR-H3 and CDR-L2 loops show less defined clustering (Fig. 2i, Fig. 2l) as is expected due to the high structural variability of CDR-H3 [27] and short length of CDR-L2 under the IMGT definition [24, 25, 28]. Visual inspection of individual clusters confirms that these embeddings capture distinct structural forms found among the samples (Fig. 2n-q). Furthermore, the predicted uncertainties output by ImmuneBuilder for the generated and held out sequences are identically distributed, with the same length-dependence being observed across both sets (Fig. S6). These results indicate that AbBFN2 covers the structural space of CDR loops that is observed in natural sequences.

### Germline gene usage

Having shown that AbBFN2 is able to accurately model CDR loops, we turned our attention to a more holistic view of antibody sequences. V(D)J recombination is the mechanism that underpins naive antibody formation, meaning that a model that has learned to reason over antibody sequences must also capture the relationships between different germline *V* (variable), *D* (diversity), and *J* (joining) genes and how they are preferentially rearranged, either explicitly or implicitly. Although V(D)J recombination is a stochastic process, certain gene pairings are overrepresented compared to others [12, 29]. We found that AbBFN2 faithfully captures these conditional distributions by comparing the pairing propensities of different gene families both within the V_H_ and V_L_ domains, as well as between them (Fig. 2r-t, Fig. S7), finding deviations of at most 1 % (IGKV1 paired with IGKJ4). This shows that at the genetic level, the biases that are present during V(D)J recombination are learned by AbBFN2, indicating that reasoning across genetic lineages and associated sequences is possible.

Since most positions remain unmutated even in a mature antibody, a bias towards these germline residues exists in many antibody-specific language models [30], yet when designing antibodies, the aim is often to move away from the germline sequence. We find that AbBFN2 can generate both naïve and mature antibody sequences, with the percent sequence identities of generations to assigned germline genes capturing the natural distribution (Fig. S8). Coupled with an ability to correctly predict the germline distance of a given sequence (as discussed in the sequence annotation task), this suggests AbBFN2 is a able to reason over the somatic divergence of generations.

### Therapeutic Antibody Profiler (TAP) Scores

Therapeutic antibodies are subject to environments that are not experienced by natural antibodies, and must therefore have biophysical properties that allow for stability under these conditions [23]. TAP uses structural models of antibodies to calculate metrics that are associated with developability: PSH (hydrophobicity), PPC (positive charge density), PNC (negative charge density), and SFvCSP (charge imbalance across the V_H_ and V_L_ domains). Sequences generated by AbBFN2 mirror the natural distributions of these metrics (Fig. S9) (including ranges that are associated with clinical-stage therapeutics), indicating that the model is able to generate therapeutic-like sequences. We note, however, that the values reported for the AbBFN2-generated samples are those predicted by the model itself, rather than the ground-truth values for the generated sequences (ground truth here refers to properties calculated for each sequence using the tool that generated the annotation for training). The accuracy of these predictions is assessed as part of the sequence annotation task.

### AbBFN2 does not memorise its training data

In order to ensure that AbBFN2 is not simply memorising the data, we calculate the distribution of minimum Hamming distances between each of the generated sequences and the training sequences (Fig. S10, Fig. 2u). We find that the generated sequences are similarly novel and diverse with respect to the training data as a held out set of natural sequences.

### AbBFN2 as a sequence annotation tool

Having examined the marginal distributions of samples generated by AbBFN2, we asked whether the model can capture the conditional dependencies between sequence data and the other labels that are included. This can be approached in two ways: either by predicting labels given the sequence of an antibody (sequence annotation), or by generating sequences subject to constraints encoded as labels (sequence design). We first examine sequence annotation and use AbBFN2 to generate all other data modalities given the full paired sequence (Fig. 1c), as described in the Methods section. In addition to accurate annotation being a valuable standalone task, successfully modeling the relationships between sequence and metadata modalities is a prerequisite for realising related inverse problems such as predicting sequences with specific germline lineages, germline sequence identities, and good developability profiles.

### Genetic lineage

The genetic lineage of an antibody encodes crucial information about structure [16, 31], and ultimately, function. Indeed, biases in germline usage in the repertoire are observed in immune responses mounted upon antigenic insult, suggesting functional differences between different genes [12, 32, 33]. Therefore, accurate identification of the germline genes and mutational distance from the corresponding naïve sequence underpin many antibody design and maturation workflows. Table 1 compares AbBFN2’s ability to predict these genetic properties from the amino acid sequence of an antibody. On every prediction task, AbBFN2 outperforms the widely used alignment-based tools ANARCI [34] and IgBLASTp [21].

**Table 1.** Performance on sequence labelling tasks. For regression, the metrics are Pearson’s R and root mean squared error (RMSE). For classification, the metric is the balanced F1 score. Missing entries indicate data modalities not predicted by a given method. Germline gene identity ground truths are the nucleotide-level values derived from OAS, assigned using IgBLASTn and as such, are more accurate than predictions made at the amino acid level. AbBFN2 is trained on directly these values but has no access to nucleotide sequences. Species and gene prediction with IgBLASTp was performed by comparing the top E-values for *V/J*-genes given human, mouse, and rat reference libraries. For Humatch and AntiBERTy, species predictions are evaluated only on species modelled by each method (human and non-human for Humatch, human and mouse for AntiBERTy). Missing entries in the table correspond to data modalities not predicted by a given method.

| Regression Tasks |  |  |  |  |  |  |
| --- | --- | --- | --- | --- | --- | --- |
| Label | Pearson’s R |  |  | RMSE |  |  |
|  | AbBFN2 | ANARCI | IgBLASTp | AbBFN2 | ANARCI | IgBLAST |
| HV gene identity (%) | **0.97** | **0.97** | **0.97** | **0.87** | 3.67 | 3.40 |
| HD gene identity (%) | **0.40** |  |  | **4.38** |  |  |
| HJ gene identity (%) | **0.80** | 0.56 | 0.57 | **1.47** | 7.19 | 7.55 |
| LV gene identity (%) | **0.97** | 0.91 | 0.96 | **0.67** | 2.81 | 2.62 |
| LJ gene identity (%) | **0.69** | 0.51 | 0.67 | **1.50** | 7.38 | 4.18 |
| PSH | **0.74** |  |  | **13.55** |  |  |
| PPC | **0.91** |  |  | **0.25** |  |  |
| PNC | **0.92** |  |  | **0.27** |  |  |
| SFvCSP | **0.90** |  |  | **2.67** |  |  |
| Classification Tasks |  |  |  |  |  |  |
| Label | AbBFN2 | ANARCI | IgBLASTp | Humatch | AntiBERTy |  |
| HV gene family | **1.00** | **1.00** | **1.00** | **1.00** |  |  |
| HD gene family | **0.68** |  |  |  |  |  |
| LV gene family | **1.00** | 0.99 | **1.00** | **1.00** |  |  |
| HV gene | **0.98** | 0.97 | 0.93 |  |  |  |
| HD gene | **0.58** |  |  |  |  |  |
| HJ gene | **0.98** | 0.88 | 0.89 |  |  |  |
| LV gene | **0.99** | **0.99** | **0.99** |  |  |  |
| LJ gene | **0.94** | 0.91 | 0.92 |  |  |  |
| LC locus | **1.00** | **1.00** | **1.00** | **1.00** |  |  |
| Species | **1.00** |  | **1.00** | **1.00 (0.92)** | **1.00 (0.93)** |  |
| PSH flag | **0.84** |  |  |  |  |  |
| PPC flag | **0.96** |  |  |  |  |  |
| PNC flag | **0.99** |  |  |  |  |  |
| SFvCSP flag | **0.97** |  |  |  |  |  |

Accurate prediction the germline gene identities given a sequence suggests that the model is able to assess to what extent a sequence has diverged from its naive clone. In the germline gene prediction task, the largest improvement is observed in the case of the V_H_ *J-*gene. We hypothesize that this is due to two contributing factors: (i) AbBFN2 being able to capture more complex statistical dependencies when compared to other tools due to its larger number of learnable parameters, and (ii) its ability to also use the whole antibody sequence to predict the V_H_ *J-*gene. Considering the fact that the residues corresponding to *J-*genes make extensive contacts with the other chain in a functional antibody, we argue that having this knowledge allows the model to reason over a richer set of dependencies. Moreover, AbBFN2 is able to predict the V_H_ *D-*gene, a task that is not tackled by currently available methods at the amino acid level due to its complexity. Owing to its very short length, the *D-*gene segment is highly ambiguous. For instance, IGHD1-7*01, IGHD1-14*01 and IGHD1-20*01 all translate to the same amino acid sequence (`GITGT`) when considering the first reading frame [35], but have differences in the underlying nucleotide sequence, making their distinction at the amino acid level challenging. Despite this, we find that AbBFN2 performs reasonably well on predicting the specific *D-*gene and better still when predicting the *D-*gene family.

### TAP developability metrics

We next asked whether AbBFN2 is able to predict the TAP flags (specific ranges of TAP scores within which clinical-stage therapeutic antibodies fall [5, 23]) for a given sequence, and found that these labels were predicted accurately by the model (Table 1). The corresponding ground truth labels are calculated using the structure of an F_V_ fragment, and the ability to AbBFN2 to reason on these properties shows an emergent implicit understanding of the three-dimensional structure of an antibody. Even the PSH score, which is known to be highly sensitive to small structural fluctuations [5], is predicted with sufficient accuracy to provide meaningful feedback about developability. Additionally, we find a similar level of performance when predicting the continuous-valued TAP metric values (Table 1), from which the flags are derived.

### AbBFN2 is able to redesign V_H_-V_L_ interfaces with sequence inpainting

Antibody language models are often evaluated on their ability to generate the full antibody given a partial sequence, also referred to as sequence inpainting (Fig. 1d). The capability to recover the remainder of the antibody demonstrates the model’s ability to reason over the whole sequence and consider inter-residue dependencies. The most straightforward metric for this is the extent to which the original sequence is recovered, amino acid recovery (AAR). We find that AbBFN2 is able to inpaint CDR loops given partial sequences and find expected trends in line with the sequence diversity of each loop. We also find that additional conditioning on germline genes increases the recovery of CDR sequences, with the most pronounced increase in the recovery of CDR-H3, highlighting the ability of AbBFN2 to reason over a joint distribution of both sequence and metadata (Fig. 3a). We present and discuss detailed results of the sequence inpainting task as well as comparisons to antibody language models in Appendix F.2. However, we argue that while easily computed, the use of AAR of inpainted CDR loop sequences is a reductive approach since multiple sequences can present similar functional profiles, as indicated by the observation that distant sequences and germline lineages can present near-identical functional paratopes [36]. Further, the evolution of an antibody sequence *in vivo* occurs in the context and presence of some antigen, which is often not considered during CDR inpainting tasks. As an alternate task to assess the ability for partial sequence generation that is functionally relevant and has less dependence on antigenic pressure, we examine interface residues of natural antibodies as inpainting targets.

**Figure 3.**
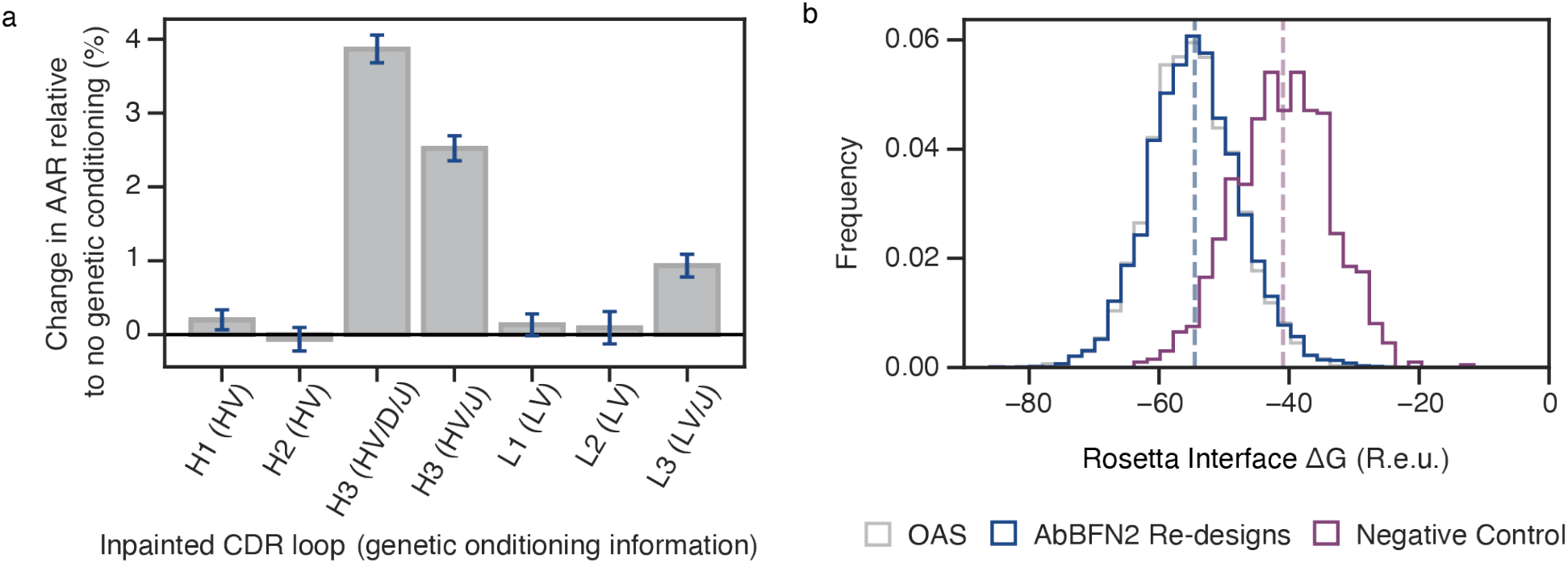
Generation of full sequences conditioned on partial sequence. **(a)** depicts the change in average amino acid recovery (AAR) rates for each CDR loop, when additional conditioning information in the form of germline genes is used apart from the remainder of the sequence and the original loop length. Each bar depicts the difference in AAR relative to inpainting experiments for the given loop where only the partial sequence and the original loop length is provided. Error bars represent the standard error of the mean. Bar heights are averages of all tested sequences (5000). We discarded any sequences where the inpainted loop does not match the original loop in length. **(b)** depicts distributions of interface stabilities of antibodies calculated by Rosetta. We predicted interface residues of hold-out sequences (grey) given the remainder of the sequence (navy) and calculated interface energies using Rosetta. As a negative control to test if Rosetta can differentiate between badly paired sequences, we randomly paired V_H_H sequences derived from OAS with V_L_ sequences from OAS. Dashed lines represent sample means. Interface residues are chosen using the same definition used by Marks *et al*. [37]

The interactions that are formed between the the V_H_ and V_L_ domains of the F_V_ region contribute heabily to the overall stability of an antibody [38], and these regions are often mutated during affinity maturation [39–41]. When inpainting these positions, we observe an AAR rate of 75.7 %. Upon examining the stability of inpainted and original sequences using Rosetta, we found no statistically significant difference (Student’s *t* = 1.21, *p* = 0.23) in the interface ΔG distributions of inpainted and original sequences, indicating that the sequences proposed by AbBFN2 are likely to be functionally valid (Fig. 3b) in terms of their pairing stability. This is in line with the observation that antibody-specific inverse folding models are often able to find alternate sequences (low AAR rates) that fold into identical structures (low RMSD). For example, it has been shown that an increase from 35 % to 60 % CDR loop AAR for inverse folding models results in a comparatively small decrease in the associated RMSD (1.03 Å to 0.95 Å) [42]. Considering the fact that, structurally, frameworks are more conserved than CDR loops, we hypothesise that a similar trend may hold true for more highly mutated positions of the framework regions, such as the interface.

### Sequence humanisation

Next, we examined the use of AbBFN2 in the context of a common antibody design workflow: sequence humanisation (Fig. 1e, Fig. 4a). This refers to the introduction of mutations to framework regions (or grafting of the candidate CDR sequences onto human frameworks) to reduce the risk of an anti-drug antibody (ADA) response being mounted in patients upon administration [43].

**Figure 4.**
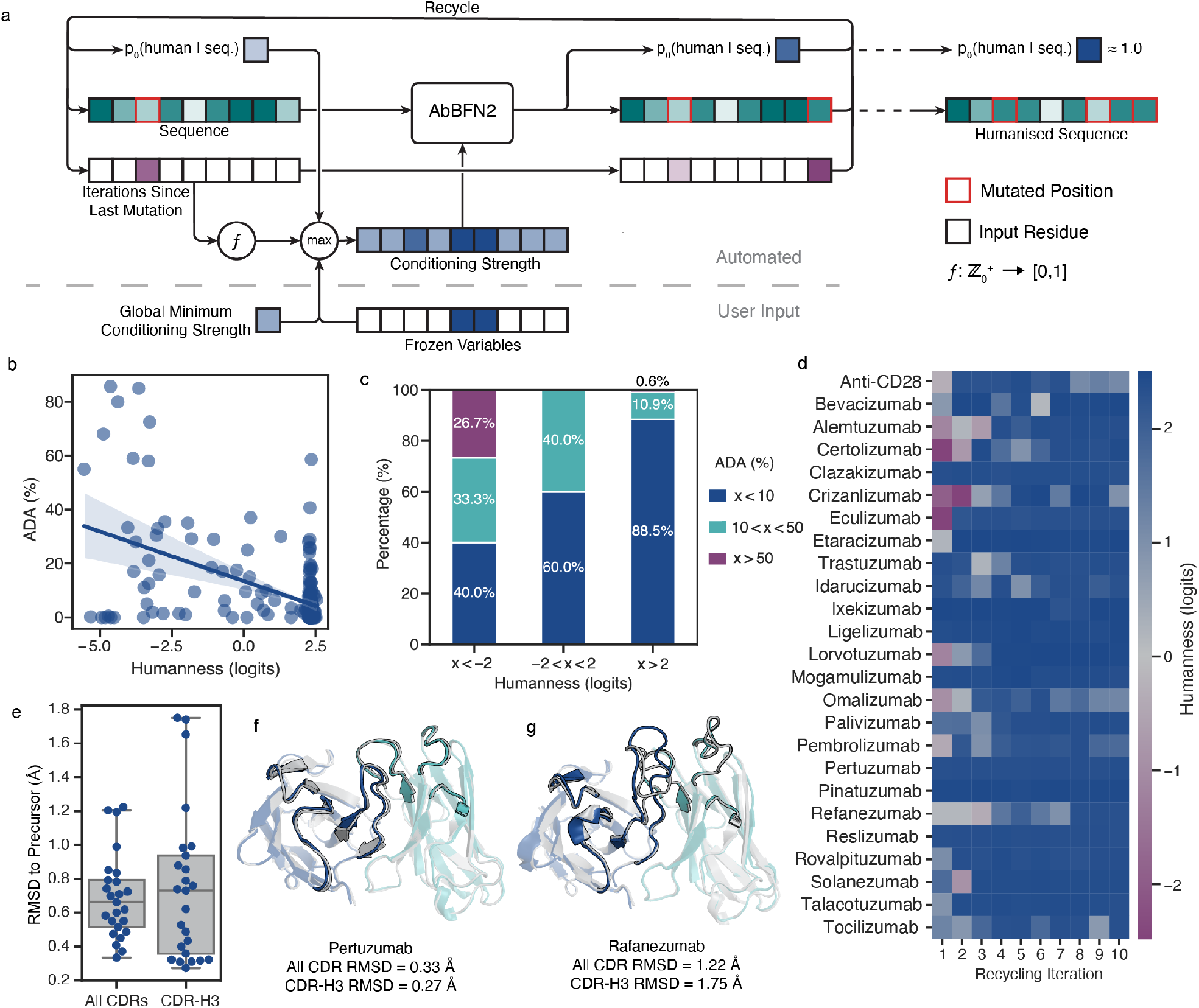
Sequence humanisation using AbBFN2. **(a)** depicts the humanisation workflow used and described in the Methods section. At each recycling step, AbBFN2 is provided with the complete current sequence. This sequence is then masked by using a combination of (i) a user-defined array specifying which positions should be frozen, (ii) a user defined minimum conditioning strength to control the overall influence of the input sequence, (iii) a buffer that keeps track of whether a position has been mutated recently, and (iv) the likelihood of the current sequence being human under the model. A lower conditioning strength of a variable translates to less influence of that variable on the sampling process, with 0 referring to a completely masked variable and 1 referring to a visible variable. **(b)** shows the correlation between the logits of the human class and the immunogenicty (percentage of trial participants developing anti-drug antibodies [ADA]) of a sequence in the Marks *et al*. data set [37]. The shaded area around the line of best fit corresponds to the 95% confidence interval calculated using bootstrapping. **(c)** shows the distributions of high-, medium- and low-immunogenicty antibodies from the Marks *et al*. data set when the human species label logits for a sequence under AbBFN2 are used to classify the antibody. **(d)** shows the humanisation trajectories of 25 clinical stage therapeutic antibodies derived from non-human precursors. The x-axis refers to the recyling iteration step, and the hue of each cell shows the human class logits for the designed sequence at each step. **(e)** shows the distribution of total CDR and CDR-H3 RMSD of redesigned sequences in (d). The first version of each sequence with *p_θ_*(species = human sequence) *≥*0.95 was extracted and folded using ImmuneBuilder. The CDRs were then aligned to ImmuneBuilder models of respective precursor sequences, and RMSDs calculated. Boxplots show the 25%, 50% (median), and 75% quartiles. Whiskers extend to a maximum of 150% of the interquartile range. **(f**,**g)** depict ImmuneBuilder models of the highest and lowest total CDR RMSD, respectively. The precursor model is shown in grey and the V_H_ and V_L_ domains of the final design are shown in navy and cyan, respectively.

### Anti-drug antibody response prediction

AbBFN2 is able to accurately predict the species of different antibody sequences (Table 1). Moreover, when considering the 211 clinical-stage therapeutic sequences collated by Marks *et al*. [37], we find that the species logits of AbBFN2 are correlated with the observed ADA responses in clinical trials (Fig. 4b). Concretely, the strength of this correlation (Pearson’s *R* = *−*0.52, *p* < 10^−15^) is in line with state-of-the-art results obtained by the autoregressive antibody language model, p-IgGen [44] (Pearson’s *R* = *−*0.53). We also found that the logit values can be used to classify candidate sequences into high-, medium-, and low-immunogenicity risk classes (Fig. 4c), similar to other humanness prediction tools [45].

### Sequence humanisation

Sequence humanisation is typically performed on a precursor sequence whose overall properties should be retained as much as possible, meaning that modifications to existing framework regions are often preferred to *de novo* design of new framework sequences [14]. Indeed, experimental humanisation often involves rounds of mutations and back-mutations, following this paradigm [46]. We developed a sampling method to recapitulate this iterative approach, illustrated in Fig. 4a and described in the Methods section. This enables AbBFN2 to run an *in silico* humanisation campaign end-to-end by repeatedly reasoning over its previous predictions and proposing candidate mutations.^1^ A sequence is considered to be successfully humanised once AbBFN2 has a > 95 % confidence in labelling it as `human`(Fig. 4d). Following the approach taken by other humanisation tools, we choose to use AbBFN2 as its own discriminator of humanness, as it has been shown that different humanisation tools learn to humanise antibodies using different sets of mutations, limiting their applicability for cross-comparison [45]. Moreover, as already seen, AbBFN2 matches or exceeds leading methods in humanness and ADA prediction.

As test cases, we chose 25 clinical-stage therapeutic antibodies derived from non-human precursors, since experimentally humanised variants exist for these sequences. AbBFN2 was able to humanise all 25 precursor sequences. In the majority of cases, AbBFN2 achieves this within a single recycling step, though some cases require up to five iterations (Fig. 4d, Fig. S12) suggesting that some input sequences present more difficult starting points. Further, the number of total mutations tends to plateau as the number of iterations increases (Fig. S11), with the average mutation count of 46.8 *±* 11.7 being comparable to the number of experimentally introduced mutations (45.1 *±* 9.7). These mutations were spread similarly across different IMGT positions in both experimental and AbBFN2-humanised sequences (Fig. S13), highlighting that AbBFN2 is able to identify the same important regions to target during humanisation campaigns. Finally, we examined the change in structure for the humanised and precursor sequences when folded using ImmuneBuilder, and found the average CDR RMSD to be (0.69 *±*0.25) Å (CDR-H3: (0.75 *±*0.44) Å) (Fig. 4e-g), indicating that the introduced mutations are likely to be well-tolerated with regard to the paratope conformation, which directly influences the ability to bind the antigen.

### Case study: Concurrent Humanisation and Removal of Sequence Liabilities Using AbBFN2

Development liability removal and humanisation are routine workflows in many antibody design campaigns, but are often performed one after the other, even though in reality antibody design is a multi-objective optimisation problem [15, 49]. This sequential approach can result in instances where previous changes are either undone, or suboptimal changes have to be picked in subsequent steps, because a position has already been mutated by a previous step in the pipeline. This can lead to more mutations having to be introduced than are necessary, or in the candidate having to be subjected to a previous step in the pipeline a second time. Here, we show the ability of AbBFN2 to optimise a sequence for two goals concurrently: humanness and TAP development liabilities (Fig. 1e).

### Optimisation procedure

We considered 91 candidates sequences that have non-human origin, at least one TAP liability and were not seen by the model during training. We again use our multi-round humanisation algorithm, adding the TAP flags as additional conditioning information (Fig. 4a) and leveraging the stochastic generation of AbBFN2 to sample four candidate solutions for each precursor sequence. We considered an optimisation trajectory successful when the sequence, once folded by using ImmuneBuilder and annotated using TAP, only had green flags (Fig. 5a-d), and when the confidence that the sequence is labeled as `human`under the model is at least 95 %.

**Figure 5.**
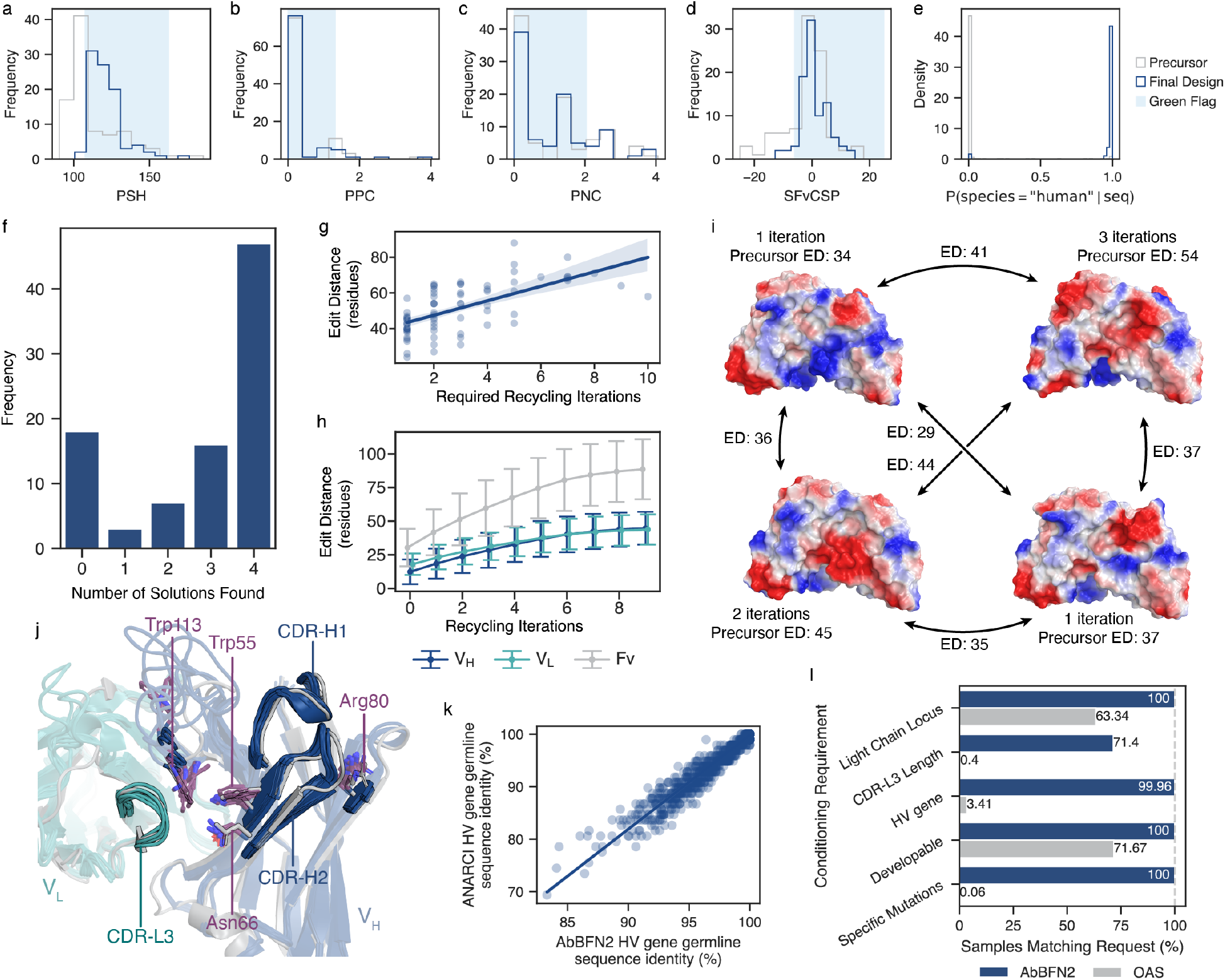
Multi-objective design using AbBFN2. **(a-i)** show results for concurrent humanisation and development liability removal from a set of 91 non-human paired sequences with at least one TAP liability in the hold-out set. For each sequence, four distinct recycling trajectories were generated, each with a total budget of 10 iterations. **(a-d)** show the precursor and final design TAP values, with the light blue region in each plot corresponding to the range of values that would be assigned a green flag. For each of the 91 precursors, we report the first instance where as many objectives as possible are fulfilled. If multiple repeats resulted in valid solutions where all objectives were satisfied, we report the one with the lowest edit distance to the precursor. **(e)** shows the precursor and design human class probability for the same samples. **(f)** depicts the number of valid solutions that were found for each of the 91 precursors across four repeats. **(g)** shows the edit distance for the first valid solution across the four optimisation trajectories per precursor. **(h)** shows the mean edit distance to the precursor sequence across all recycling trajectories, with error bars representing standard deviations. **(i)** shows the four valid solutions for one precursor, with the edit distance (ED) to the precursor and the required number of iterations. Arrows between the sequences are marked with edit distances between the solutions. Each solution is folded using ImmuneBuilder and coloured according to electrostatic potential using Pymol. **(j-l)** show results for conditional generation of VRC01-like antibodies. **(j)** shows alignments of folded AbBFN2-generated samples and VRC01 (PDB: 3NGB), with residues and regions of interest labelled. **(k)** shows the distribution of V_H_ *V*-gene identities across generated samples. The x-axis corresponds to predictions directly output by AbBFN2, whereas the y-axis shows the calculated identities using ANARCI. The line of best fit is shown, with shaded areas corresponding to the 95% confidence interval calculated using bootstrapping. **(l)** shows comparisons of how frequent the requested features are in the held-out OAS data and the conditionally generated AbBFN2 samples.

### Optimised Sequences

AbBFN2 is able to successfully optimise most sequences (Fig. 5a-f) within the first iteration of the recycling workflow, with a maximum of 10 iterations required and an average of 56.9 mutations used (Fig. 5g, Fig. 5h).^2^ We also find that independent trajectories given the same starting sequence result in diverse sequences being generated. For a randomly selected sequence, visualisation of the resultant structures also shows different surface properties (Fig. 5i), confirming that the desired objectives can be satisfied in a number of ways.

In total, 18 of the 91 of the starting sequences were not successfully optimised within the alloted compute budget (Fig. 5f). While we expect that further independent trajectories may lead to optimised solutions for some of these sequences, we found that the failing objective in 15 was an unchanged TAP PNC (negative charge density) or PPC (positive charge density) metric (Fig. 5c). The CDRs of these sequences—which were kept constant during the optimisation procedure—had more charged residues than those for which optimised solutions were identified (Fig. S14). We therefore suggest that this may mean that the CDRs themselves already present an antibody that is highly unlikely to be developable. Indeed, a number of our starting sequences appear to be more difficult than others, as for some of these starting points, only a subset of the four repeats produce succesfully optimised candidates. Ultimately, we cannot confirm if these 18 sequences are fundametally undevelopable, required more iterations for valid optimisation in line with some of the other starting points, or are outside of the model capabilities. Regardless, this is in line with the fact that initial candidate selection, whether automated or by a human expert, can have an appreciable impact on downstream design.

Taken together, these results suggest that AbBFN2 can be used as a sequence redesign tool to optimise multiple attributes of an antibody in a concerted manner, provided that the optimisation goals do not present a paradoxical set of conditions.

### Case study: Generation of a developable library of rare antibodies

Having tested AbBFN2’s ability to redesign a given sequence with multiple target objectives, we next sought to understand its ability to generate antibody sequences *de novo* while respecting multiple objectives (Fig. 1f, Fig. 5j-l) by generating a library of VRC-01-like antibodies. VRC-01 is an unusual anti-HIV antibody [50], with a combination of multiple rare characteristics that are not found in the germline repertoires of most people. As such, both natural and display-based induction of this class of antibody is challenging. Namely, these antibodies are marked by (i) a human *V-*gene lineage of IGHV1-2, (ii) a *κ* locus light chain, (iii) a CDR-L3 loop length of 5 residues, and (iv) position-specific residues on the V_H_ framework (Trp55, Asn66, Arg80) and CDR-H3 loop (Trp113) [51].

We generate 2500 sequences with AbBFN2, conditioned both to match the VRC-01 properties above, and to satisfy our developability metrics (all TAP flags must be of class `green`). The generated library is then analysed by calculating the empirical CDR lengths, checking the set sequence positions, and computing TAP metrics from structures obtained with ImmuneBuilder (Fig. 5j); ultimately concluding that 1715 sequences satisfy all conditioning properties. It is interesting to compare this highly-tailored generation of AbBFN2 with traditional pLM-based methods, where adaptation to a specific target distribution is typically achieved by fine-tuning on a representative dataset. Whilst AbBFN2 does not require such fine-tuning, in highly specific cases—such as VRC-01—it may not even be a viable option due to the scarcity of the desired properties in natural repetoires. Concretely, within the *∼* 2 M paired chains in OAS, and despite approximately 10 000 sequences coming from repertoires of HIV-infected individuals, only 21 VRC-01-like antibodies exist that have all desired characteristics in combination.

Satisfying the requested characteristics is essential for effective library generation, however it is also necessary that relationships between variables that have not been conditioned on are appropriately shifted based on the conditioning information. For instance, when conditioning on a specific V_H_ *V-*gene, the CDR-H1 and CDR-H2 sequences should reflect the underlying conditional distribution. To this end, we examined a number of dependencies that might be influenced by the conditioning information, in-depth results for which we present in Appendix appendix F.3. Briefly, we find that (i) the spread of germline sequence identities to the IGHV1-2 germline sequence (Fig. 5k) matches the expected diversity (Fig. S8), (ii) that the V_L_ *V-*gene family distribution of the generated samples is consistent with that of natural V_L_ sequences with short CDR-L3 loops and *κ* lineages, and (iii) that the diversity of V_H_ CDR loop sequences is in line with the diversity expected when restricting the V_H_ *V*-gene (specifically, we find low diversity in CDR-H1/2—which only have sequence contributions from this gene—and high diversity in CDR-H3, which also has contributions from the *D/J*-genes).

Taken together, these results suggest that AbBFN2 allows for finely tunable *de novo* generation of antibodies with an arbitrary set of user-requested parameters (Fig. 5l). Moreover, AbBFN2 is able to explore diversity within the portion of sequence space that the conditioning information restricts it to. We note that we use VRC-01-like antibodies to demonstrate that AbBFN2 is able to generate sequences with combinations of rare attributes, and argue that if a similarly generated library were to be used for screening, additional conditioning on the CDR-H3 sequences of VRC-01-like antibodies would be likely to increase success rates.

## Discussion

Here, we present AbBFN2, a multi-modal foundation model for paired antibody sequences that is able to faithfully capture sequences in its training distribution, along with relevant metadata. Owing to its training scheme, AbBFN2 is able to unconditionally generate natural-like sequences *de novo*, but also allows for conditional generation of sequences with an arbitrary set of conditioning parameters. We also demonstrate the ability of the model to accurately label sequences with genetic attributes and potential development liabilities, as well as inpaint parts of a sequence given the remainder. Further, AbBFN2 can be used for antibody design, optimising sequences whilst paying attention to multiple objectives, compressing a traditional computational design pipeline into a single step. This has the advantage of minimising the risk of a later step undoing changes made by an earlier one.

Given the flexibility of AbBFN2, we note that the way the model is queried by the user can have considerable impacts on the output samples. For instance, we found that humanisation required multiple recycling iterations, akin to chain-of-thought approaches used by large language models [52]. Similar approaches have previously been used in computational antibody design [53]. We also found that human expertise is necessary when interacting with the model to avoid paradoxical prompting, as shown in our evaluation of liability removal in antibodies with highly charged CDR loops. We argue that as such multi-modal, flexible approaches become more commonplace in generative artificial intelligence for biological applications, prompt engineering considerations will also become necessary in order to correctly and sensibly steer the generative process.

AbBFN2 covers a breadth of sequence metadata relating to sequence, genetics, and developability, but is agnostic on the antigen, epitope, and paratope information as well as structural details of the antibody. As BFNs are naturally well-adapted to dealing with both discrete (sequence, gene labels, etc.) and continuous (structural coordinates, germline sequence identity, etc.) data, a natural extension of AbBFN2 would be to include further data modes including the structure of the antibody. Further, although paired antibody sequence data are becoming more commonplace, the inclusion of unpaired data will allow for a more extensive set of dependencies to be captured by the model.

## Data Availability

Training, validation, and test data are deposited in Zenodo [54] under https://doi.org/10.5281/zenodo.15237125, including metadata that were derived in-house. All data used in the study were taken from the publicly available Observed Antibody Space [19, 20] database.

## Code Availability

The inference code for AbBFN2 can be found on https://github.com/instadeepai/AbBFN2. A web application is provided under https://abbfn2.labs.deepchain.bio. Both the code and web application are provided under a noncommercial license.

## Acknowledgements

This work was supported by Cloud TPUs from Google’s TPU Research Cloud (TRC). We thank Miles McGibbon and Benoit Gaujac for feedback on the manuscript.

## Author Contributions

BG and TB conceived the research. BG curated training and validation data sets, trained the model, developed and implemented sampling methods, wrote code, performed experiments, and analysed data. MB prepared open-source code for release. TB and TA wrote code and assisted work with Bayesian Flow Networks. SC supported the development of soft inpainting for Bayesian Flow Networks. LC supported the design of experiments. AG developed and implemented sampling methods for conditional generation. AL supported the interpretation and presentation of results. BG and TB wrote the manuscript. TB supervised the project.

## Competing Interests

The authors declare no competing interests.

## Methods

### Algorithmic details

BFNs [17] aim to learn joint probability distribution of *N* variables – which we denote as *p*(*x*) = *p*(*x*_1_, …, *x_N_*) – by modelling the parameters of a distribution over the data – *θ* = [*θ*_1_, …, *θ_N_*] where *p*(*x_i_*|*θ_i_*) is the probability distribution of the *i*-th variable. The training process of a BFN can be framed as a noisy communication protocol; where the network is initially fully uninformed about the ground truth sample, *x*, and has to learn to predict it given a series of corrupted observations. Intuitively, the fewer bits of information the BFN requires to accurately predict the ground truth, the better the model has learned the underlying distribution. Full details on the model architecture, and how this is formalized into a training objective and practically implemented is provided in Supplementary Information A and B.

At inference time, the BFN is used to transition *θ* from a fully uninformed initial prior to a coherent distribution over the data from which a sample can be drawn. Moreover, as well as unconditional generation of all variables, this sampling process can be modified to access conditional distributions, *p*(*x*∉*x*_cond_| *x*_cond_), where arbitrary subsets of the variables set to desired values, *x*_cond_. Specifically, in this work we consider score-based guidance [55] and particle filtering [56] for conditional generation tasks. Full details on sampling methodologies are again provided in Supplementary Information C; with the sampling algorithms themselves provided in Supplementary Information Algorithms 16, 23 and 24.

### Data

#### Sequence Numbering and Structural Modelling

All sequences were numbered according to the IMGT scheme [28] using ANARCI [34] (CDR1: IMGT residues 27-38, CDR2: IMGT residues 56-65, CDR3: IMGT residues 105-117, Framework: IMGT residues 1-128 and not within CDR definitions). We structurally modelled F_V_ regions using the ABodyBuilder2 model within the ImmuneBuilder Suite [22] using provided pre-trained weights.

#### Therapeutic Antibody Profiler (TAP)

To calculate development liabilities, we used an in-house implementation of the Therapeutic Antibody Profiler [23]. TAP calculates four metrics: (i) PSH (patches of hydrophobic residues), (ii) PPC (patches of positively charged residues), (iii) PNC (patches of negatively charged residues), and (iv) SFvCSP (V_H_/V_L_ charge symmetry). These values are then compared to known clinical-stage therapeutics (CSTs), and percentile scores are used to assign a flag for each property. A green flag corresponds to values of a metric that is commonly observed in CSTs, an amber flag corresponds to values which are rarely observed in CSTs, and red flags correspond to values outside of the known CST distribution. To account for changes due to differences in how solvent-accessible surface areas are calculated, we also recalibrated our in-house implementation using the sequences used to calibrate the latest version of TAP [5].

#### Observed Antibody Space (OAS)

We downloaded paired sequences from the Observed Antibody Space (OAS) [19, 20] on May 24^th^, 2024. We removed sequences based on whether they had fewer than 8 residues missing from either terminus of the variable domain whether they were productive sequences as per the OAS IgBLASTn [21] annotation and whether they had missing conserved cysteine residues, yielding a total of 2,031,524 paired V_H_ and V_L_ sequences.

### Sequence Clustering

We clustered the described sequences by following the clustering method used for AbLang2 [30]. Namely, CDR-3 sequences are extracted from each sequence and a unique cluster is created for each unique CDR-3 sequence. Within each cluster, the whole chain is clustered using Linclust [57] at a sequence identity threshold of 95% and `-cov-mode 1`(where there are overlapping clusters within a CDR-3 cluster, these are handled internally by Linclust). The final cluster for a variable domain is then determined by its whole sequence cluster within the given CDR-3 loop cluster. Finally, the heavy and light chains are paired. Two antibodies are therefore considered to be in the same cluster if they have the same CDR-H3 and CDR-L3 loops, and if both the V_H_ and V_L_ domains show sequence similarity greater than 95 %. We constructed the train, test, and hold-out sets by selecting 99 %, 0.5 %, and 0.5 % of the unique final clusters, respectively.

### Data Modes

AbBFN2 is trained on a total of 45 “data modes”. 14 of these come from the V_H_ and V_L_ sequences. Namely, each sequence is broken up into three CDR loops, and padded at the centre of the loop to the longest length observed for that loop in the processed OAS data, ensuring that the processed sequences follow the IMGT scheme [28], where insertions are placed at the centre of the loop. The four framework regions of each chain are similarly processed but padding tokens appended to the sequence. Start and end tokens are added to each sequence data mode. Another six data modes were used to categorically encode the lengths of the CDR loops. We found that encoding these as categorical variables yielded more faithful conditional generation results than encoding them as discretised variables.

Ten data modes are assigned to the germline genes of the antibody as annotated in OAS using IgBLASTn [21]. Three of these represent the heavy chain *V-, D-* and *J-*genes, with another three representing the families of these genes (i.e. IGHV1-2 and IGHV1, respectively). The remaining four data modes are assigned to the the light chain *V-* and *J-*genes in an analogous manner. We did not include the alleles of these genes, as we found that the relatively small amount of training data available was not enough to capture allelic variants effectively. These data modes are encoded as categorical variables. Another five data modes represent the percentage sequence identity to each assigned germline gene, encoded as a continuous variable.

One data mode was used for the species label found in paired OAS (human, mouse, and rat antibodies), and another data mode represents the light chain locus (*κ* and *λ*). Both were encoded as categorical variables. The last eight data modes represent the TAP values: four continuous variables representing the PSH, PPC, PNC, and SFvCSP metrics, and four discrete variables representing the assigned flag for each (green, amber, and red).

For each of the categorical variables, we also included an additional class referring to unknown values. Vales that appeared fewer than 100 times in the training set, as well as missing data were assigned this category.

We note that this set of features has some level of redundancy (for instance, the PSH score and corresponding flag). We still include these features as the provide an easier way to interact with the model during conditional generation tasks. For instance, a user might be interested in generating antibodies with developable PSH values (green flag), but might not necessarily want to restrict the generative process to a specific value of the PSH metric within this space. Similarly, a user might want to generate antibodies coming from a specific germline gene family without explicitly requesting a specific gene.

### Training Scheme

We trained AbBFN2 using a batch size of 4096 for 30 000 steps. We used the Adam optimiser [58] with *β*_1_ = 0.9 and *β*_2_ = 0.98. We initialised training with a learning rate of 0, which was then increased linearly to 5 × 10^−4^ over 1000 steps, after which it is kept constant, and clipped gradient norms to 1. During sampling, we adopted the same method used to train ProtBFN [18], where the probability of selecting a cluster in the training set is proportional to the square root of its size. Once a cluster is selected, a member from within that cluster is then selected at random. This debiases the training regime away from large clusters while ensuring that variability within each cluster can be observed by the model during training. Since the loss per data mode is calculated independently, we found that a smaller number of total steps provided balance between overfitting on easy to learn data modes and underfitting on more difficult data modes.

### Sequence Design via Sample Recycling and Soft Inpainting

We found that attempting to redesign an existing antibody for some target property was challenging in a single iteration. For instance, during sequence humanisation with specific CDR sequences, we would sample conditioned on the CDR loops and the human label for the species data mode, but found that AbBFN2 would often generate samples belonging to the species of the parental antibody. Further, by only conditioning on the CDRs, any information pertaining to the precursor framework regions is lost when a common aim in antibody design workflows is to maintain as much of the parental sequence as possible. To address these issues, we developed an alternate sampling technique consisting of a series of recycling steps wherein the model is queried using its own output from a previous iteration. Further, during each sampling iteration, the model is conditioned on the requested attributes (CDR loops, species label, etc.) throughout the generative process, but is also “softly” conditioned on the parental framework sequence by exposing the parental sequence during the first *n*% of the sampling steps, in a manner analogous to differential diffusion [59]. For instance, a soft conditioning of strength 0.5 results in the framework being visible to the model in the first half of the sampling process. A conditioning strength of 0 equates to complete masking of the variable, whereas a conditioning strength of 1 refers to traditional conditional generation as outlined in Section C.2.

Since the conditioning strength can be set per variable, each residue can in theory contribute to the conditioning information in a variable manner. In our case, we set the per-residue conditioning strength using the following scheme. Each CDR loop residue had a constant conditioning strength of 1. The conditioning strength for all other residues was set to the probability under AbBFN2 that the species label is `human`given the whole antibody sequence, *p_θ_*(`human`| **x**) where **x** is the antibody sequence, clipped to a minimum value of 0.2 (this requires an initial pass of the sequence through the model ahead of the first iteration). We found the clipping to be necessary to prevent the conditioning strength being set to ≈ 0 (meaning the model pays no attention to the framework) when the sequence is classed as non-human. The output sequence of the previous iteration, **x***^′^* is used as input for the next step, with the framework sequence conditioning strength being updated according to *p_θ_*(`human`**x***^′^*), also derived from the output of the previous iteration. Intuitively, this approach ensures that when a sequence is non-human (*p_θ_*(`human`| **x***^′^*) is low) AbBFN2 is able to make more mutations on the framework due to the low conditioning strength. As a sequence becomes more human (*p_θ_*(`human`| **x***^′^*) is high), the conditioning strength increases, thereby preventing the model from making large changes. When a sequence is human (*p_θ_*(`human`| **x***^′^*) *≈* 1), the sequence cannot change anymore, and the final solution is therefore “locked in”.

While the above approach eventually converges to a valid human sequence, we observed frequent back-mutations, wherein AbBFN2 reverts mutations it has already introduced. These reversion events greatly increase the number of iterations required. To mitigate this effect and encourage the model to commit to previously made mutations, we modified the conditioning strength for positions that have already been mutated. However, to allow the model to eventually revise these positions again, we decay this value as a mutation becomes older. Specifically, if a position was mutated *n* iterations ago, *c*(*n*) is given by

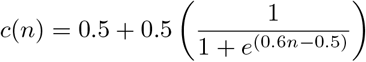

This ratcheting mechanism ensures drift away from the starting sequence whilst still allowing eventual revision of previously edited positions. Lastly, *c*(*n*) is clipped to be at least *p_θ_*(`human`|**x***^′^*).

In order to ensure that the solution also converges to a single *V-*gene during sequence humanisation (since although technically human, chimeric antibodies containing sequences coming from multiple *V-*genes are not natural), we used the germline-likeness score proposed by Chinery *et al*. [45] to first introduce seed mutations, with the closest human germline being inferred using IgBLASTp [21].

### Validation

#### Sequence Similarity Calculations

To calculate sequence similarity across antibodies, we first aligned sequences to match the most common 200 IMGT positions as described by Olsen *et al*. [60] This ensures that functionally similar residues are aligned with each other and also standardises the length of each sequence. These length-matched sequences were then compared using Hamming distances, as implemented in `scipy`[61].

#### Structural Similarity Calculations using Dynamic Time Warping

In order to compare the structures of CDR loops, we first structurally modelled sequences using ABodyBuilder2 [22], part of the ImmuneBuilder suite. We then compared loops using dynamic time warping (DTW) (Algorithm 27). DTW allows for the comparison of loops of different lengths, similar to how the Needlemann-Wunsch algorithm [62] allows comparison across sequences with different sizes by finding the optimal alignment path through a cost matrix. When two loops are of identical length and the main diagonal is used to traverse the cost matrix, it reduces down to a conventional RMSD calculation. All calculations are performed on the backbone C, C*α*, and N residues. For each CDR loop we first performed a pairwise alignment of all loops of interest on their corresponding flanking anchor residues (five residues on either side of the loop) and calculated the pairwise anchor RMSD values. The medoid across this step was then chosen as the reference, and all other loops were aligned on the anchors of this structure. If a structure had an anchor RMSD of > 1.5 Å, it was removed, as good anchor aligbnment is required for DTW to produce appropriate comparisons. We then calculated the pairwise CDR backbone DTW distances for all remaining, anchor-aligned structures using the implementation in `DTAIDistance`[63], and generated t-SNE embeddings for the generated pairwise distance matrix using `scikit-learn`[64].

#### Interface Stability Calculations

To assess the interface stability of a V_H_-V_L_ pair, we first structurally modelled the pair using ABodyBuilder2. We then used `pyrosetta`[65] to perform relaxation using the FastRelax protocol [66] with default settings and five rounds of repacking and relaxation. The V_H_-V_L_ interface analysis was then performed using the InterfaceAnalyzerMover module of `pyrosetta`, recoding the interface Δ*G* value reported.

#### Germline gene matching using IgBLASTp and ANARCI

To perform germline gene matching using IgBLASTp, we used default settings, with germline libraries built using the IMGT database [28] on Sep 27, 2024. Where species information is available, we used the correct species library. Where species information was not available, we calculated the most likely germline genes across all three modelled species (human, mouse, and rat) and picked the alignment with the lowest E-score. To perform germline gene matching using ANARCI, we used default settings, limiting the set of species for alignment to human, mouse, and rat.

#### AntiBERTy, AbLang2, and Humatch

For sequence inpainting tasks, AntiBERTy [67] and AbLang2 [30] were used with default settings. For AbLang2, both the V_H_ and V_L_ sequences were provided. Humatch [45] was run using settings and thresholds as described in the tutorial example of the released code. A sequence was classed as human if it had a probability of at least 0.5 for any of the human *V*-gene family categories.

## Footnotes

1 We also note that similar inference time compute scaling approaches have been shown to improve the output of both autoregressive [47], and diffusion-based [48] generative models.

2 In total, the optimisation of all 91 sequences with a compute budget of 4 repeats at 10 iterations each takes 2.5 hours on a v4-8 TPU instance.

## Notes

https://github.com/instadeepai/AbBFN2

https://abbfn2.labs.deepchain.bio

https://doi.org/10.5281/zenodo.15237125

### Competing Interest Statement

The authors have declared no competing interest.

