## Supplementary Information for "AbBFN2: A flexible antibody foundation model based on Bayesian Flow Networks"

**D Auxiliary Algorithms** 13

**E Supplementary Figures** 15

**F Supplementary Results** 25

#### List of Algorithms

1 Forward pass of the output network (Φ). . . . 5
2 Encode input distribution discrete. . . . 5
3 Encode input distribution continuous. . . . 5
4 Regression head . . . 6
5 Decode output distribution discrete . . . 6
6 Decode output distribution continuous . . . 6
7 Training loss . . . 7
8 Noise schedule discrete . . . 7

© 2025 InstaDeep. All rights reserved

9 Noise schedule continuous . . . 8
10 Sample flow distribution discrete . . . 8
11 Sample flow distribution continuous . . . 8
12 Residual loss discrete . . . 8
13 Residual loss continuous . . . 8
14 Reconstruction loss discrete . . . 9
15 Reconstruction loss continuous . . . 9
16 Unconditional sampling . . . 9
17 Prior distribution discrete . . . 9
18 Prior distribution continuous . . . 10
19 Sample receiver distribution discrete . . . 10
20 Sample receiver distribution continuous . . . 10
21 Update distribution discrete . . . 10
22 Update distribution continuous . . . 10
23 SDE sampling . . . 11
24 Twisted SDE sampling . . . 12
25 Particle logit discrete . . . 13
26 Particle logit continuous . . . 13
27 Dynamic Time Warping . . . 14

#### List of Figures

S1 Position-specific amino acid propensities across the hold-out set. . . . 15
S2 Position-specific amino acid propensities across generated samples. . . . 16
S3 Logo plots of position specific residue propensity differences. . . . 17
S4 Heatmaps of position specific residue propensity differences. . . . 18
S5 Average differences in amino acid frequencies across loops. . . . 19
S6 Relationships between sequence length and ImmuneBuilder predicted error scores. . . . 20
S7 Frequencies of gene pairings and differences across generated and hold-out samples. . . . 21
S8 Distributions of sequence identities to germline genes. . . . 21
S9 Distributions of Therapeutic Antibody Profiler (TAP) metrics. . . . 22
S10 Minimum Hamming distances of the held out and generated samples to the training set. . . . 22
S11 Number of total mutations accumulated during sequence humanisation. . . . 22
S12 Number of mutations and steps required for sequence humanisation. . . . 23
S13 Position-specific mutations in experimentally and computationally humanised sequences. . . . 24
S14 Charges of CDR residues in successful and unsuccessful optimisation trajectories. . . . 25

### A. Model architecture

#### A.1. Data modes

AbBFN2 models 45 *data modes*, each of which is comprised one or more related variables. We highlight that each data mode may be comprised of continuous or discrete variables, summarised in Tables S1 and S2, respectively. Note that by convention, a specific data mode does not mix variable modalities to allow for modality-specific encoder and decoder blocks as discussed in appendix A.2.

| Mode Name | Description | Num vars. | Type | Range |
|---|---|---|---|---|
| $V_H$ *V*-gene ID | % ID of $V_H$ *V*-gene to germline | 1 | continuous | $[64.0, 100.0]$ |
| $V_H$ *D*-gene ID | % ID of $V_H$ *D*-gene to germline | 1 | continuous | $[74.0, 100.0]$ |
| $V_H$ *J*-gene ID | % ID of $V_H$ *J*-gene to germline | 1 | continuous | $[74.0, 100.0]$ |
| $V_L$ *V*-gene ID | % ID of $V_L$ *V*-gene to germline | 1 | continuous | $[66.0, 100.0]$ |
| $V_L$ *J*-gene ID | % ID of $V_L$ *J*-gene to germline | 1 | continuous | $[77.0, 100.0]$ |
| PSH | TAP surface hydrophobicity score | 1 | continuous | $[72.0, 300.0]$ |
| PPC | TAP patches of positive charge score | 1 | continuous | $[0.0, 7.5]$ |
| PNC | TAP patches of negative charge score | 1 | continuous | $[0.0, 10.0]$ |
| SFvCSP | TAP charge asymmetry score | 1 | continuous | $[-55.0, 55.0]$ |

**Table S1 | Overview of continuous data modes included in AbBFN2.** For continuous data modes, we clipped data to the reported ranges before linearly scaling each range to $[-1, 1]$. For germline identity data modes, missing values were replaced with $100.0$. For TAP metrics, missing values were replaced with the median of each distribution.

As a Bayesian Flow Network [1], AbBFN2 does not model raw data directly, but instead models a distribution over the data. Concretely, this means that the inputs and outputs to the network are per-variable parameters of this distribution, $\theta_i$ such that $p(x_i|\theta_i)$ is the probability distribution of the $i$-th variable. The exact form of $\theta_i$ depends on whether the variable is continuous or discrete:

- **Continuous variables** are modelled with a Gaussian distribution parametrized with a mean, $\mu$, and variance, $\rho$. Therefore, in this case $\theta_i = \{\mu_i, \rho_i\}$ such that $p(x_i|\theta_i) = \mathcal{N}(x_i|\mu_i, \rho_i)$. Following the original formulation of Graves *et al.* [1], the variance of the output distribution is set to match that of the input - therefore $\rho_i$ is not modelled explictly by the network.
- **Discrete variables** are modeled as a categorical distribution parameterized by a set of logits, and so for $K$ classes have $\theta_i = [z_i^{(1)}, \ldots, z_i^{(K)}]$ such that $p(x_i = k|\theta_i) = \frac{\exp(z_i^{(k)})}{\sum_{k'=1}^{K} \exp(z_i^{(k')})}$.

We henceforth denote the parameters of all variables modelled by AbBFN2 as $\boldsymbol{\theta}$, the set of parameters that comprise the $i$-th data mode as $\boldsymbol{\theta}^{(\text{dm-i})}$ and the parameters of the $j$-th variable as $\theta_j$. The specific implementation of functions can differ depending on whether the variables to which they are applied are continuous or discrete. As a notional convenience, we will assume that a function for which both a continuous and discrete implementation is provided can be called on arbitrary sets of parameters, with the mapping to the correct modality-specific implementation assumed implicitly.

| Mode Name | Description | Num vars. | Type | Num classes. ($K$) |
|---|---|---|---|---|
| $V_H$-FR1 | Seq. of $V_H$ framework 1 | 43 | discrete | 32 |
| $V_H$-FR2 | Seq. of $V_H$ framework 2 | 32 | discrete | 32 |
| $V_H$-FR3 | Seq. of $V_H$ framework 3 | 60 | discrete | 32 |
| $V_H$-FR4 | Seq. of $V_H$ framework 4 | 14 | discrete | 32 |
| $V_H$-CDR1 | Seq. of CDR-H1 loop | 24 | discrete | 32 |
| $V_H$-CDR2 | Seq. of CDR-H2 loop | 27 | discrete | 32 |
| $V_H$-CDR3 | Seq. of CDR-H3 loop | 60 | discrete | 32 |
| $V_L$-FR1 | Seq. of $V_L$ framework 1 | 38 | discrete | 32 |
| $V_L$-FR2 | Seq. of $V_L$ framework 2 | 29 | discrete | 32 |
| $V_L$-FR3 | Seq. of $V_L$ framework 3 | 50 | discrete | 32 |
| $V_L$-FR4 | Seq. of $V_L$ framework 4 | 15 | discrete | 32 |
| $V_L$-CDR1 | Seq. of CDR-L1 loop | 22 | discrete | 32 |
| $V_L$-CDR2 | Seq. of CDR-L2 loop | 18 | discrete | 32 |
| $V_L$-CDR3 | Seq. of CDR-L3 loop | 29 | discrete | 32 |
| $V_H$ *V*-gene | Name of $V_H$ *V*-gene | 1 | discrete | 201 |
| $V_H$ *D*-gene | Name of $V_H$ *D*-gene | 1 | discrete | 49 |
| $V_H$ *J*-gene | Name of $V_H$ *J*-gene | 1 | discrete | 7 |
| $V_H$ *V*-gene fam. | Name of $V_H$ *V*-gene family | 1 | discrete | 10 |
| $V_H$ *D*-gene fam. | Name of $V_H$ *D*-gene family | 1 | discrete | 8 |
| $V_H$ *J*-gene fam. | Name of $V_H$ *J*-gene family | 1 | discrete | 7 |
| $V_L$ *V*-gene | Name of $V_L$ *V*-gene | 1 | discrete | 220 |
| $V_L$ *J*-gene | Name of $V_L$ *J*-gene | 1 | discrete | 13 |
| $V_L$ *V*-gene fam. | Name of $V_L$ *V*-gene family | 1 | discrete | 18 |
| $V_L$ *J*-gene fam. | Name of $V_L$ *J*-gene family | 1 | discrete | 13 |
| $V_L$ LC locus | Locus of the light chain | 1 | discrete | 3 |
| Species | Species of the sequence | 1 | discrete | 4 |
| $V_H$-CDR1 length | Length of CDR-H1 loop | 1 | discrete | 13 |
| $V_H$-CDR2 length | Length of CDR-H2 loop | 1 | discrete | 13 |
| $V_H$-CDR3 length | Length of CDR-H3 loop | 1 | discrete | 33 |
| $V_L$-CDR1 length | Length of CDR-L1 loop | 1 | discrete | 12 |
| $V_L$-CDR2 length | Length of CDR-L2 loop | 1 | discrete | 5 |
| $V_L$-CDR3 length | Length of CDR-L3 loop | 1 | discrete | 15 |
| PSH flag | Developability flag for PSH score | 1 | discrete | 4 |
| PPC flag | Developability flag for PPC score | 1 | discrete | 4 |
| PNC flag | Developability flag for PNC score | 1 | discrete | 4 |
| SFvCSP flag | Developability flag for SFvCSP score | 1 | discrete | 4 |

**Table S2 | Overview of data modes included in AbBFN2.** For discrete data modes, the number of classes including an additional class for unknown values are provided.

#### A.2. Output network

The output network transforms the *input distribution*, $p_{\mathrm{i}}(\boldsymbol{x}|\boldsymbol{\theta})$ to a predicted *output distribution*, $p_{\mathrm{o}}(\boldsymbol{x}|\hat{\boldsymbol{\theta}})$ where $\hat{\boldsymbol{\theta}} = \text{outputNetwork}(\boldsymbol{\theta}, \dots)$. The overall architecture consists of an encoder and decoder module for each data mode; and a shared backbone. A forward pass of the network is described in algorithm 1.

**Algorithm 1** Forward pass of the output network ($\Phi$).

**def** outputNetwork( $\boldsymbol{\theta} = [\boldsymbol{\theta}^{(\text{dm-1})}, \boldsymbol{\theta}^{(\text{dm-2})} \dots]$, cumulative accuracy $\beta$, hidden dimension $d_\text{h} = 1280$, feedforward dimension $d_\text{ffn} = 1280$, $N_\text{layers} = 33$, $N_\text{head} = 20$ ):

*# Encode input distribution parameters*

**for** $i = \text{dm-1}, \text{dm-2}, \dots$ **do**

$\quad \boldsymbol{h}^{(i)} = \text{encode}^{(i)}(\boldsymbol{\theta}^{(i)}, d_\text{h})$ ▷ $\boldsymbol{h}^{(i)} \in \mathbb{R}^{\text{len}(\boldsymbol{\theta}^{(i)}) \times d_\text{h}}$

*# Process through shared backbone network*

$\boldsymbol{h} = [\boldsymbol{h}^{(1)}, \boldsymbol{h}^{(2)}, \dots]$

**for** $i = 1$ to $N_\text{layers}$ **do**

$\quad \boldsymbol{h} \mathrel{+}= \text{MultiHeadAttention}(\text{RMSNorm}(\boldsymbol{h}), N_\text{head})$

$\quad \boldsymbol{h} \mathrel{+}= \text{FeedForward}(\text{RMSNorm}(\boldsymbol{h}), d_\text{ffn})$

*# Decode output distribution parameters*

$[\boldsymbol{h}^{(1)}, \boldsymbol{h}^{(2)}, \dots] = \boldsymbol{h}$

**for** $i = \text{dm-1}, \text{dm-2}, \dots$ **do**

$\quad \boldsymbol{\theta}^{(i)} = \text{decode}^{(i)}(\boldsymbol{h}^{(i)}, \boldsymbol{\theta}^{(i)}, \beta)$

**return** $\boldsymbol{\theta} = [\boldsymbol{\theta}^{(\text{dm-1})}, \boldsymbol{\theta}^{(\text{dm-2})} \dots]$

**Encoders** The the per-data mode encoders project the input parameters for each variable to a shared embedding space. In place of a time encoding (as used in [1] and traditional diffusion-like architectures), AbBFN2 instead uses an entropy encoding as proposed in [2]. For continuous variables, this choice is arbitrary as the entropy of the input distribution is a deterministic function of time as given by the noise schedule. However, the entropy of the input distribution for discrete variables is not deterministic, and it has been proposed that this can lead to a mismatch between the entropy levels at a fixed time during the training and sampling procedure. Algorithm 2 and algorithm 3 detail the encoding functions for a single discrete or continuous variable, respectively.

**Algorithm 2** Encode input distribution discrete.

**def** encode( $\theta = [z^{(1)}, \dots, z^{(K)}]\}$, hidden dimension $d_\text{h}$, ):

$\boldsymbol{p} = [p^{(1)}, \dots, p^{(K)}] = \text{softmax}(\theta)$

$\boldsymbol{h} = \text{Linear}(\boldsymbol{p})$ ▷ $\boldsymbol{h} \in \mathbb{R}^{d_\text{h}}$

*# Apply entropy encoding*

$\tilde{H} = -\frac{1}{\log K} \sum_{i=1}^{K} p^{(i)} \log p^{(i)}$ ▷ Normalized entropy

$\boldsymbol{h} \mathrel{+}= \text{Linear}(\tilde{H})$

**return** $\boldsymbol{h}$

**Algorithm 3** Encode input distribution continuous.

**def** encode( $\theta = \{\mu, \rho\}$, hidden dimension $d_\text{h}$, ):

$\boldsymbol{h} = \text{Linear}(\mu)$ ▷ $\boldsymbol{h} \in \mathbb{R}^{d_\text{h}}$

*# Apply entropy encoding*

$\tilde{H} = \text{sigmoid}\left(\frac{1}{2} \log\left(\frac{2\pi}{\rho}\right) + \frac{1}{2}\right)$ ▷ Normalized entropy

$\boldsymbol{h} \mathrel{+}= \text{Linear}(\tilde{H})$

**return** $\boldsymbol{h}$

**Backbone** The main trunk of the network is a standard Transformer architecture, comprising stacked residual multi-head attention (MHA) and feed-forward (FFN) blocks. To promote stable training, the final linear layer in each residual block, as well as the query projection matrix in the MHA blocks, is zero-initialized. The MHA blocks use rotary positional embeddings [3] with a frequency coefficient (commonly denoted by $\theta$ in prior work, but this should not be confused with our definition of $\theta$ as distribution parameters) of $10\,000$. The FFN blocks employ SwiGLU activations, and to maintain the same parameter count as standard (non-gated) FFN layers with dimension $d_{\text{ffn}}$, the linear projections in these gated blocks effectively operate on a $\frac{2}{3}d_{\text{ffn}}$ hidden dimension.

**Decoders** The per-data mode decoders transform the backbone embeddings to predictions of the output distribution parameters. The main computational unit is a small FFN as defined in algorithm 4. For discrete variables, this unit directly predicts logits of the output distribution; however, for continuous variables we follow Graves et al. [1] and reparameterise the output distribution as a noise prediction task. All decoder units clip output distribution parameters to a maximum magnitude for numerical stability. Note that all continuous data modes are rescaled such that $\mu \in [-1, 1]$. Algorithm 5 and algorithm 6 detail the decoding functions for a single discrete or continuous variable, respectively.

**Algorithm 4** Regression head

**def** regressionHead( $\boldsymbol{h} \in \mathbb{R}^{d_\text{h}}$, output dimension $d_\text{o}$ ):
$\boldsymbol{h} = \text{Linear}(\text{LayerNorm}(\boldsymbol{h}))$ ▷ $\rightarrow \boldsymbol{h} \in \mathbb{R}^{2 \cdot d_\text{h}}$
$\boldsymbol{h} = \text{gelu}(\boldsymbol{h})$
$\boldsymbol{h} = \text{Linear}(\text{LayerNorm}(\boldsymbol{h}))$ ▷ $\rightarrow \boldsymbol{h} \in \mathbb{R}^{d_\text{o}}$
**return** $\boldsymbol{h}$

**Algorithm 5** Decode output distribution discrete

**def** decode( $\boldsymbol{h} \in \mathbb{R}^{d_\text{h}}$, ~~$\theta = [z^{(1)}, \ldots, z^{(K)}]$, $\beta$,~~ max logit magnitude $z_{\text{max}} = 10$ ):
$\boldsymbol{z} = \text{regressionHead}(\boldsymbol{h}, d_\text{o} = K)$ ▷ $\boldsymbol{z} \in \mathbb{R}^K$
$\boldsymbol{z} = \text{clip}\left(\boldsymbol{z}, -z_{\text{max}}, z_{\text{max}}\right)$
**return** $\theta = \boldsymbol{z}$

**Algorithm 6** Decode output distribution continuous

**def** decode( $\boldsymbol{h} \in \mathbb{R}^{d_\text{h}}$, $\theta = \{\mu, \rho\}$, $\beta$, output limits $\mu_{\text{lim}} = 1$ ):
$\varepsilon = \text{regressionHead}(\boldsymbol{h}, d_\text{o} = 1)$ ▷ $\varepsilon \in \mathbb{R}$
$\gamma = \text{clip}(\frac{\beta}{1+\beta}, \text{min} = 1\text{e}{-9})$ ▷ Clipped for numerical stability.
$\mu = \frac{\mu}{\gamma} - \sqrt{\frac{1-\gamma}{\gamma}}\varepsilon$
$\mu = \text{clip}(\mu, -\mu_{\text{lim}}, \mu_{\text{lim}})$
**return** $\theta = \{\mu, \rho = 1 + \beta\}$

#### B. Training

The training process can be framed as a noisy communication protocol; where the BFN is initially fully uninformed about the ground truth sample, $\boldsymbol{x}$, and then receives a series of observations $\{\boldsymbol{y}^{(i)}\}_{i \in \{0,1\}}$ with which it needs to increasingly refine it's belief over the value of $\boldsymbol{x}$. At each step $i$, we have:

1. **Noisy observation.** A noisy version of the original data, $\boldsymbol{y}^{(i)}$, is sampled from the *sender distribution* $p_\text{s}(\boldsymbol{y}^{(i)} | \boldsymbol{x}, \alpha^{(i)})$ where $\alpha^{(i)}$ is an accuracy parameter determining how informative (i.e. how noisy) the sample is
2. **Bayesian inference.** All observed noisy signals, $\{\boldsymbol{y}^{(i)}, \ldots, \boldsymbol{y}^{(i)}\}$, are combined to obtain a set of parameters $\boldsymbol{\theta}^{(i)}$ via Bayesian inference on a per-variable basis. These parameters form the *input distribution*.
3. **Neural refinement.** The parameters $\boldsymbol{\theta}^{(i)}$ are passed to the output network, which predicts a refined *output distribution*, $p_\text{o}^{(i)}$. This allows the prediction for each variable to be refined using information contained in other variables.

4. **Prediction of the next noisy observation.** The output distribution $p_{\mathrm{o}}^{(i)}$ is then subjected to the same noise process as the ground-truth data, yielding a *receiver distribution* $p_{\mathrm{r}}^{(i+1)}$, which represents the model's prediction for the next noisy observation.

Although the above process is described as a discrete step process, in practice it can also be expressed in continuous time, where information and probability mass change smoothly for $t = 0 \to t = 1$, hence the name *Bayesian Flow*. By doing so, we can consider the distribution of input parameters obtained as $t$ to be the Bayesian aggregation of all information obtained from the sender distribution up to that point, and call this the *flow distribution*, $p_{\mathrm{f}}^{(i)}$. The training objective for the BFN is to minimise the total number of nats required to transmit the data; which is given by the sum of two terms; the residual, $\mathcal{L}_{\mathrm{res}}$, and reconstruction, $\mathcal{L}_{\mathrm{recon}}$, losses (detailed in algorithms 12 to 15).

Algorithm 7 details the calculation of the BFN training loss. Further details on specific steps are provided below, and for a full derivation and discussion of the BFN algorithm the reader is reffered to the original work of Graves *et al.* [1].

**Algorithm 7** Training loss

**def** loss( $\Phi$, $\boldsymbol{x}$, noise parameters $\boldsymbol{\eta} = \{\beta_j^{(1)}$ or $\sigma_j^{(1)}\}_{j=1,2,\dots}$ ):

*# Compute residual loss*

$t \sim U[0, 1]$

$\alpha^{(t)}, \beta^{(t)} =$ noiseSchedule$(t, \boldsymbol{\eta})$

$\boldsymbol{\theta}^{(t)} =$ sampleFlow$(\boldsymbol{x}, \beta^{(t)})$

$\hat{\boldsymbol{\theta}}^{(t)} =$ outputNetwork$(\boldsymbol{\theta}^{(t)}, \beta^{(t)})$

$\mathcal{L}_{\mathrm{res}} =$ residualLoss$(\boldsymbol{x}, \hat{\boldsymbol{\theta}}^{(t)}, \alpha^{(t)})$

*# Compute reconstruction loss.*

$\_, \beta^{(1)} =$ noiseSchedule$(1, \boldsymbol{\eta})$

$\boldsymbol{\theta}^{(1)} =$ sampleFlow$(\boldsymbol{x}, \beta^{(1)})$

$\hat{\boldsymbol{\theta}}^{(1)} = \Phi(\boldsymbol{\theta}^{(1)})$

$\mathcal{L}_{\mathrm{recon}} =$ reconstructionLoss$(\boldsymbol{x}, \hat{\boldsymbol{\theta}}^{(1)}, \boldsymbol{\eta})$

**return** $\mathcal{L}_{\mathrm{res}} + \mathcal{L}_{\mathrm{recon}}$

**Noise schedules** The noise applied to each variable is a fixed function of time, with the information contained in the sender distribution samples at time $t$ being given by the aforementioned accuracy parameter, $\alpha^{(t)}$. The accuracy schedule is then the accumulated accuracy up to $t$, $\beta^{(t)} = \int_{t'=0}^{t} \alpha^{(t)}$. Importantly, $\beta^{(t)}$ increases monotonically from $\beta^{(0)} = 0$, i.e. fully uninformative samples, to $\beta^{(1)}$, which is fixed to set the desired accuracy over the entire process.

Concretely, algorithm 8 gives the noise schedule for discrete variables – which takes the form of the heuristic schedule proposed in Graves et al. [1]. When training AbBFN2, we use $\beta^{(1)} = 1.0$ (corresponding to $\sigma^{(1)} = 1/\sqrt{2} \approx 0.71$) for all discrete variables. For continuous variables (algorithm 9), the functional forms are chosen such that the entropy of the input distribution decreases linearly with $t$. Note that in this case it is more intuitive to set the accuracy of the process by fixing $\sigma_1$ to the standard deviation of the input distribution at $t = 1$, hence the functions are provided in these terms. In the case of AbBFN2, we set $\sigma^{(1)} = 1/\sqrt{5001} \approx 0.0141$ (corresponding to $\beta^{(1)} = 5000.0$) for all continuous variables.

**Algorithm 8** Noise schedule discrete

**def** noiseSchedule($t$, total accuracy $\beta^{(1)} = 1.0$):

$\alpha = 2t\beta^{(1)}$

$\beta = t^2\beta^{(1)}$

**return** $\alpha, \beta$

**Algorithm 9** Noise schedule continuous

**def** noiseSchedule($t$, final noise $\sigma^{(1)} = 0.0141$):
$\alpha = -\frac{2 \ln \sigma^{(1)}}{(\sigma^{(1)})^{2t}}$
$\beta = (\sigma^{(1)})^{-2t} - 1$
**return** $\alpha, \beta$

**Flow distribution** Algorithms 10 and 11 show how to sample flow distribution for a single variable.

**Algorithm 10** Sample flow distribution discrete

**def** sampleFlow($x$, $\beta$):
$e_x = \text{oneHot}(x, \texttt{num\_classes} = K)$
$\boldsymbol{z} \sim \mathcal{N}\left(\beta(Ke_x - 1), \beta K \mathbb{I}\right)$ ▷ Sample logits
**return** $\theta = \boldsymbol{z}$

**Algorithm 11** Sample flow distribution continuous

**def** sampleFlow($x$, $\beta$):
$\gamma = \frac{\beta}{1+\beta}$
$\mu \sim \mathcal{N}\left(\gamma x, \gamma(1-\gamma)\right)$
$\rho = 1 + \beta$
**return** $\theta = \{\mu, \rho\}$

**Residual loss** The residual loss is the number of nats required to transmit the sender samples $\{\boldsymbol{y}^{(t)}\}_{t=0}^{t=1}$. Given the nats to transmit a sample from sender, $p_{\mathrm{s}}$, to receiver, $p_{\mathrm{r}}$ is $D_{\mathrm{KL}}(p_{\mathrm{s}} \parallel p_{\mathrm{r}})$, it follows that the overall residual loss is given by

$$\mathcal{L}_{\mathrm{res}} = \mathop{\mathbb{E}}_{t \sim U[0,1]} D_{\mathrm{KL}}(p_{\mathrm{s}}^{(t)} \parallel p_{\mathrm{r}}^{(t)}). \tag{1}$$

Practically, algorithm 12 and algorithm 13 show how to calculate residual losses for a single discrete and continuous variable, respectively.

**Algorithm 12** Residual loss discrete

**def** residualLoss( $\theta$=$[z^{(1)}, \ldots, z^{(K)}]\}$, $x$, $\alpha$ ):
$e_x = \text{oneHot}(x, \texttt{num\_classes} = K)$
$\hat{e}_x = \text{softmax}(\theta)$
**return** $K\frac{\alpha}{2}\sum_{k=1}^{K}(e_x^{(k)} - \hat{e}_x^{(k)})^2$

**Algorithm 13** Residual loss continuous

**def** residualLoss( $\theta$=$\{\mu, \rho\}$, $x$, $\alpha$ ):
**return** $\frac{\alpha}{2}(\mu - x)^2$

**Reconstruction loss** The reconstruction loss is the number of nats required to transmit $\boldsymbol{x}$ at $t = 1$ (i.e. accounting for any remaining mismatch between the BFN belief and ground truth after all sender samples are processed). This is given by

$$\mathcal{L}_{\mathrm{recon}} = -\mathop{\mathbb{E}}_{p_{\mathrm{f}}(\boldsymbol{\theta}|\boldsymbol{x}, t=1)} \ln p_{\mathrm{o}}(\boldsymbol{x}|\boldsymbol{\theta}; t = 1). \tag{2}$$

We note that continuous data requires infinite precision to reconstruct, therefore practically it is assumed that there is some finite measurement noise $\sigma$ on $\boldsymbol{x}$. By doing so, the reconstruction loss can be defined as the KL divergence between $\mathcal{N}(\boldsymbol{x}, \sigma^2\mathbb{I})$ and $p_{\mathrm{o}}(\boldsymbol{x}|\boldsymbol{\theta}; t = 1)$. In this work, we set this measurement noise to be equal to the standard deviation of the input distribution at $t = 1$, and thus $\sigma = \sigma_1$.

Practically, algorithm 14 and algorithm 15 show how to calculate reconstruction losses for a single discrete and continuous variable, respectively.

**Algorithm 14** Reconstruction loss discrete

**def** reconstructionLoss( $\theta$=$[z^{(1)}, \dots, z^{(K)}]\}$, $x$, ~~total accuracy $\beta_T$~~ ):
**return** $-z^{(x)}$ ▷ CCE loss

**Algorithm 15** Reconstruction loss continuous

**def** reconstructionLoss( $\theta$=$\{\mu, \rho\}$, $x$, final noise $\sigma_1$ ):
**return** $\frac{1}{8\sigma_1^2}(\mu - x)^2$

As a final note, it has been observed that for a suitably parameterized schedule, learning to minimise the reconstruction loss is trivial, as the input distribution at $t = 1$ is already very close to $\boldsymbol{x}$ [1, 2]. Therefore some prior works have elected to train using only the residual loss. In this work, we do optimise the reconstruction loss but choose not to allocate half of the training compute to doing so. Instead, we practically implement a small modification to algorithm 7 where 95 % of samples are trained with the residual loss with the remaining 5 % using the reconstruction loss. A simple re-weighting of these terms allows us to recover the desired balanced training objective, $\mathcal{L}_{\text{res}} + \mathcal{L}_{\text{recon}}$, without using balanced compute for each term.

### C. Sampling

#### C.1. Unconditional generation

When unconditionally sampling from the BFN, there is no ground truth $\boldsymbol{x}$ from which to generate a sender or flow distribution. Therefore, we instead use the previous steps receiver distribution to 'hallucinate' sender samples in the next step (as the receiver distribution is the model's belief over the sender distribution). With this single change, the multi-step communication process previously discussed in appendix B to motivate the training process can be modified to generate new samples, as detailed in algorithm 16.

**Algorithm 16** Unconditional sampling

**def** sample($N_{\text{steps}}$, $\boldsymbol{\eta}$):
$\theta^{(0)} = $ priorDistribution()
$\hat{\theta}^{(0)} = $ outputNetwork($\boldsymbol{\theta}^{(0)}, \beta = 0$)
**for** $t^{(i)} = 0, \frac{1}{N_{\text{steps}}}, \dots \frac{N_{\text{steps}}-1}{N_{\text{steps}}}$ **do**
    $(_, \beta^{(i)}), (_, \beta^{(i+1)}) = $ noiseSchedule($t^{(i)}, \boldsymbol{\eta}$), noiseSchedule($t^{(i+1)}, \boldsymbol{\eta}$)
    $\alpha^{(i)} = \beta^{(i+1)} - \beta^{(i)}$
    $\boldsymbol{y}^{(i)} = $ sampleReceiver($\hat{\theta}^{(i)}, \alpha^{(i)}$)
    $\theta^{(i+1)} = $ updateDistribution($\theta^{(i)}, \boldsymbol{y}^{(i)}, \alpha^{(i)}$)
    $\hat{\theta}^{(i+1)} = $ outputNetwork($\boldsymbol{\theta}^{(i+1)}, \beta^{(i+1)}$)
**return** $\hat{\theta}^{(1)}$

**Prior distribution** The input distribution is initialised as a fully uninformed prior as described for individual discrete and continuous variables in algorithm 17 and algorithm 18, respectively.

**Algorithm 17** Prior distribution discrete

**def** priorDistribution($t$):
**return** $\theta = \mathbf{0}$ ▷ $\theta \in \mathbb{R}^K$

**Algorithm 18** Prior distribution continuous

**def** priorDistribution($t$):
1: **return** $\theta = \{\mu{=}0, \rho{=}1\}$

**Receiver distribution** As discussed above, the receiver distribution is the output distribution of the network after being subject to the same noising process as the ground truth data. The function forms for sampling a noisy observation from the receiver distribution for for a single discrete and continuous variable are provided in algorithm 19 and algorithm 20, respectively.

**Algorithm 19** Sample receiver distribution discrete

**def** sampleReceiver( $\theta{=}[z^{(1)}, \dots, z^{(K)}]\}$, $\alpha$ ):
1: $\boldsymbol{p} = \text{softmax}(\theta)$
2: $\boldsymbol{y} \sim \mathcal{N}\left(\alpha(K\boldsymbol{p} - 1),\, \alpha K \mathbb{I}\right)$ ▷ $\boldsymbol{y} \in \mathbb{R}^K$
3: **return** $\boldsymbol{y}$

**Algorithm 20** Sample receiver distribution continuous

**def** sampleReceiver( $\theta{=}\{\mu, \rho\}$, $\alpha$ ):
1: $y \sim \mathcal{N}\left(\mu,\, \alpha^{-1}\right)$ ▷ $y \in \mathbb{R}$
2: **return** $y$

**Update distribution** Given a receiver sample, $\boldsymbol{y}$, the distribution parameters of the BFN are updated to form the input distribution for the next step. This is a Bayesian update performed per-variable, and the closed-form update for both discrete and continuous variables are provided in algorithm 21 and algorithm 22, respectively.

We note that an optional score function with the same dimension as the predicted distribution parameters can also be provided. Whilst not used for unconditional generation, conditional generation algorithms described below can leverage score-based guidance which is applied at this step.

**Algorithm 21** Update distribution discrete

**def** updateDistribution( $\theta{=}[z^{(1)}, \dots, z^{(K)}]\}$, $\boldsymbol{y}$, $\alpha$, score function $\boldsymbol{s}$ = None ):
1: $\boldsymbol{z} = \theta + \boldsymbol{y}$
2: **if** $\boldsymbol{s} \neq$ None **then**
3: $\quad \boldsymbol{z} \mathrel{+}= \alpha K \boldsymbol{s}$
4: **return** $\theta = \boldsymbol{z}$

**Algorithm 22** Update distribution continuous

**def** updateDistribution( $\theta{=}\{\mu, \rho\}$, $y$, $\alpha$, score function $s$ = None ):
1: $\mu = \frac{\mu\rho + \alpha y}{\rho + \alpha}$
2: **if** $s \neq$ None **then**
3: $\quad \mu \mathrel{+}= \left(\frac{1}{\rho} - \frac{1}{\rho+\alpha}\right) s$
4: $\rho \mathrel{+}= \alpha$
5: **return** $\theta{=}\{\mu, \rho\}$

##### C.2. Score-based conditional generation

A BFN learns the joint distribution of variables $p(\boldsymbol{x}) = p(x_1, \dots, x_N)$. Given desired values for some subset of these variables $\boldsymbol{x}_c$, our conditional generation task is to sample from the distribution $p(\boldsymbol{x}|\boldsymbol{x}_c)$.

Our first algorithm to achieve this is inspired by classifier guidance as developed and popularised for conditional generation from diffusion models [4]. Score-based diffusion models [5] model the continuous-time dynamics of data subject to a Gaussian noising process, $\boldsymbol{x}^{(t)}$, by training a neural network to model the 'score' function, $\nabla \log p(\boldsymbol{x}^{(t)})$. Given conditioning data, the score function becomes $\nabla \log p(\boldsymbol{x}^{(t)}|\boldsymbol{x}_\mathrm{c})$ which can be re-expressed using Bayes' Theorem as $\nabla \log p(\boldsymbol{x}^{(t)}) + \nabla \log p(\boldsymbol{x}_\mathrm{c}|\boldsymbol{x}^{(t)})$. That is, the conditional score function becomes the sum of the unconditional score function and the gradients of a classifier determining the (log) probability of the conditioning data.

Although BFNs were not originally proposed, nor have been presented thus far, as score-based models, recent work from Xue et al. [6] has shown that the Bayesian Flow can be expressed as as stochastic differential equation (SDE) and provided closed-form score functions that govern the evolution dynamics. Whilst they leveraged this to propose faster sampling methods based on SDE and ODE formulations explored in diffusion models, we can equally extend this analysis to allow score-based conditional generation.

Concretely, instead of an external classifier, the BFN can be used to compute the probability of the conditioning data under the output distribution $p_\mathrm{o}(\boldsymbol{x}_\mathrm{c})$. Using this, we can compute the conditional modification to the score function $\boldsymbol{s} = \nabla_{\boldsymbol{x}_\mathrm{c}} \log p_\mathrm{o}(\boldsymbol{x}_\mathrm{c})$. Ultimately, this allows us to use classifier guidance in our SDE sampling function as presented in algorithm 23.

**Algorithm 23** SDE sampling

**def** sampleSDE( $N_\mathrm{steps}$, $\boldsymbol{\eta}$, conditioning data $\boldsymbol{x}_c$, conditioning mask $\boldsymbol{m}_c$, max score $s_\mathrm{max} = 1.0$, score scale $c = 1.0$ ):

$\boldsymbol{\theta}^{(0)} =$ priorDistribution()
$\hat{\boldsymbol{\theta}}^{(0)} =$ outputNetwork$(\boldsymbol{\theta}^{(0)}, \beta = 0)$
$\boldsymbol{s}^{(0)} = \mathrm{clip}\left(\nabla_{\boldsymbol{x}_c} \log p_\mathrm{o}(\boldsymbol{x}_c|\boldsymbol{\theta}^{(0)}), -s_\mathrm{max}, s_\mathrm{max}\right)$ ▷ Score function
$\boldsymbol{s}^{(0)} = c\boldsymbol{s}^{(0)}$
**for** $t^{(i)} = 0, \frac{1}{N_\mathrm{steps}}, \ldots \frac{N_\mathrm{steps}-1}{N_\mathrm{steps}}$ **do**
  $(\_, \beta^{(i)}),\ (\_, \beta^{(i+1)}) =$ noiseSchedule$(t^{(i)}, \boldsymbol{\eta})$, noiseSchedule$(t^{(i+1)}, \boldsymbol{\eta})$
  $\alpha^{(i)} = \beta^{(i+1)} - \beta^{(i)}$
  $\boldsymbol{y}^{(i)} =$ sampleReceiver$(\hat{\boldsymbol{\theta}}^{(i)}, \alpha^{(i)})$
  $\boldsymbol{\theta}^{(i+1)} =$ updateDistribution$(\boldsymbol{\theta}^{(i)}, \boldsymbol{y}^{(i)}, \alpha^{(i)}, \boldsymbol{m}_c \cdot \boldsymbol{s}^{(i)})$ ▷ Score-guided update
  $\hat{\boldsymbol{\theta}}^{(i+1)} =$ outputNetwork$(\boldsymbol{\theta}^{(i+1)}, \beta^{(i+1)})$
  $\boldsymbol{s}^{(i+1)} = \mathrm{clip}\left(\nabla_{\boldsymbol{x}_c} \log p_\mathrm{o}(\boldsymbol{x}_c|\boldsymbol{\theta}^{(i+1)}), -s_\mathrm{max}, s_\mathrm{max}\right)$ ▷ Score function
  $\boldsymbol{s}^{(i+1)} = c\boldsymbol{s}^{(i+1)}$
**return** $\hat{\boldsymbol{\theta}}^{(1)}$

##### C.3. Particle-based conditional generation

Our second conditional generation algorithm extends the score-guided SDE to further use Sequential Monte Carlo (SMC) to maintain an ensemble of weighted sampling trajectories ('particles'). We build on the work of Wu et al. [7] to 'twist' the proposal and re-weighting scheme of SMC such that the distribution of particles approaches the target conditional distribution. At a high level, this means that we run $K$ sampling trajectories using our SDE sampling scheme; but at each step resample the particles according to some particle weight, $w^{(k)}$. This algorithm, which we call twisted SDE sampling, is provided in algorithm 24.

**Algorithm 24** Twisted SDE sampling

**def** sampleTwistedSDE( $N_{\text{steps}}$, $\boldsymbol{\eta}$, conditioning data $\boldsymbol{x}_c$, conditioning mask $\boldsymbol{m}_c$, max score $s_{\max} = 1.0$, num particles $K = 8$ ):

**for** $k = 1, \ldots K$ **do**
- $\boldsymbol{\theta}_k^{(0)} = \text{priorDistribution}()$
- $\hat{\boldsymbol{\theta}}^{(0)} = \text{outputNetwork}(\boldsymbol{\theta}^{(0)}, \beta = 0)$
- $\boldsymbol{s}_k^{(0)} = \text{clip}\left(\nabla_{\boldsymbol{x}_c} \log p_{\text{o}}(\boldsymbol{x}_c | \boldsymbol{\theta}_k^{(0)}), -s_{\max}, s_{\max}\right)$
- $w_k^{(0)} = 0.0$ ▷ Initialize particle logit

**for** $t^{(i)} = 0, \frac{1}{N_{\text{steps}}}, \ldots \frac{N_{\text{steps}}-1}{N_{\text{steps}}}$ **do**
- $\left\{\boldsymbol{\theta}_k^{(i)}, \hat{\boldsymbol{\theta}}_k^{(i)}, \boldsymbol{s}_k^{(i)}\right\}_{k=1}^{K} \sim \text{Multinomial}\left(\left\{\boldsymbol{\theta}_k^{(i)}, \hat{\boldsymbol{\theta}}_k^{(i)}, \boldsymbol{s}_k^{(i)}\right\}_{k=1}^{K} \middle| \text{logits} = \left\{w_k^{(i)}\right\}_{k=1}^{K}\right)$ ▷ Resample particles
- $(\_, \beta^{(i)}),\ (\_, \beta^{(i+1)}) = \text{noiseSchedule}(t^{(i)}, \boldsymbol{\eta}),\ \text{noiseSchedule}(t^{(i+1)}, \boldsymbol{\eta})$
- $\alpha^{(i)} = \beta^{(i+1)} - \beta^{(i)}$
- **for** $k = 1, \ldots K$ **do**
  - $\boldsymbol{y}_k^{(i)} = \text{sampleReceiver}(\hat{\boldsymbol{\theta}}_k^{(i)}, \alpha^{(i)})$
  - $\boldsymbol{\theta}_k^{(i+1)} = \text{updateDistribution}(\boldsymbol{\theta}_k^{(i)}, \boldsymbol{y}_k^{(i)}, \alpha^{(i)}, \boldsymbol{m}_c \cdot \boldsymbol{s}_k^{(i)})$
  - $\hat{\boldsymbol{\theta}}_k^{(i+1)} = \text{outputNetwork}(\boldsymbol{\theta}_k^{(i+1)}, \beta^{(i+1)})$
  - $\boldsymbol{s}_k^{(i+1)} = \text{clip}\left(\nabla_{\boldsymbol{x}_c} \log p_{\text{o}}(\boldsymbol{x}_c | \boldsymbol{\theta}_k^{(i+1)}), -s_{\max}, s_{\max}\right)$
  - $w_k^{(i+1)} = \text{particleLogit}(\boldsymbol{\theta}_k^{(i)}, \boldsymbol{\theta}_k^{(i+1)}, \hat{\boldsymbol{\theta}}_k^{(i)}, \hat{\boldsymbol{\theta}}_k^{(i+1)}, \boldsymbol{y}_k^{(i)}, \alpha^{(i)}, \boldsymbol{x}_c, \boldsymbol{m}_c)$ ▷ Compute particle logit

**return** $\left\{\hat{\boldsymbol{\theta}}_k^{(1)}\right\}_{k=1}^{K}$

**Particle weights** The particle weight calculation is adapted from Algorithm 1 of [7]. Concretely, the weight of a particle is given by

$$w^{(i+1)} = \frac{p(\boldsymbol{\theta}^{(i+1)} | \boldsymbol{\theta}^{(i)})}{\tilde{p}(\boldsymbol{\theta}^{(i+1)} | \boldsymbol{\theta}^{(i)}, \boldsymbol{x}_c)} \cdot \frac{\tilde{p}(\boldsymbol{x}_c | \boldsymbol{\theta}^{(i+1)})}{\tilde{p}(\boldsymbol{x}_c | \boldsymbol{\theta}^{(i)})}, \tag{3}$$

where $p$ and $\tilde{p}$ are the ('untwisted' and 'twisted') probability distributions if the ground-truth conditioning data isn't and is used to provide score-based guidance to the distribution update between $\boldsymbol{\theta}^{(i)}$ and $\boldsymbol{\theta}^{(i+1)}$.

Now, we can note that the Bayesian update of $\boldsymbol{\theta}^{(i)}$ to $\boldsymbol{\theta}^{(i+1)}$ is a deterministic function of the receiver sample $\boldsymbol{y}$. Therefore, we can consider the probability of the receiver samples that would give rise to the observed update both without and with score guidance (which we denote $\boldsymbol{y}'$ and $\boldsymbol{y}$, respectively), to see that

$$p(\boldsymbol{\theta}^{(i+1)} | \boldsymbol{\theta}^{(i)}) = p_{\text{r}}(\boldsymbol{y}'^{(i)} | \boldsymbol{\theta}^{(i)}, \alpha^{(i)}),\ \text{where } \boldsymbol{\theta}^{(i+1)} = \text{updateDistribution}(\theta^{(i)}, \boldsymbol{y}'^{(i)}, \alpha^{(i)}, \text{None}) \tag{4}$$

and

$$\tilde{p}(\boldsymbol{\theta}^{(i+1)} | \boldsymbol{\theta}^{(i)}) = p_{\text{r}}(\boldsymbol{y}^{(i)} | \boldsymbol{\theta}^{(i)}, \alpha^{(i)}),\ \text{where } \boldsymbol{\theta}^{(i+1)} = \text{updateDistribution}(\theta^{(i)}, \boldsymbol{y}^{(i)}, \alpha^{(i)}, \boldsymbol{m}_c \cdot \boldsymbol{s}^{(i)}). \tag{5}$$

Putting equations (4) and (5) back into equation (3), and noting that $\tilde{p}(\boldsymbol{x}_c | \boldsymbol{\theta}) = p_{\text{o}}(\boldsymbol{x}_c | \boldsymbol{\theta})$ we can express, the particle logits as

$$\log w^{(i+1)} = \left[\log p_{\text{r}}(\boldsymbol{y}'^{(i)} | \boldsymbol{\theta}^{(i)}, \alpha^{(i)}) - \log p_{\text{r}}(\boldsymbol{y}^{(i)} | \boldsymbol{\theta}^{(i)}, \alpha^{(i)})\right] + \left[\log p_{\text{o}}(\boldsymbol{x}_c | \boldsymbol{\theta}^{(i+1)}) - \log p_{\text{o}}(\boldsymbol{x}_c | \boldsymbol{\theta}^{(i)})\right]. \tag{6}$$

Algorithm 25 and algorithm 26 detail the calculation of the particle logit for a single discrete and continuous variable, respectively. Note that the particle logits are computed for each variable independently, and then summed to determine a particle's overall likelihood of being re-sampled.

**Algorithm 25** Particle logit discrete

**def** particleLogit( $\theta^{(i)}$, $\theta^{(i+1)}$, $\hat{\theta}^{(i)}$, $\hat{\theta}^{(i+1)}$, $\boldsymbol{y}$, ~~$\alpha$~~, $x_c$, $m_c$ ):

$\boldsymbol{p} = \text{softmax}(\hat{\theta}^{(i)})$

$p_\text{r}^{(i)} \leftarrow \mathcal{N}\left(\alpha(K\boldsymbol{p} - 1),\, \alpha K\mathbb{I}\right)$ ▷ Receiver distribution

$p_\text{o}^{(i)},\, p_\text{o}^{(i+1)} = \text{Categorical}(\texttt{logits} = \hat{\theta}^{(i)}),\, \text{Categorical}(\texttt{logits} = \hat{\theta}^{(i+1)})$ ▷ Output distribution(s)

$\boldsymbol{y}' = \theta^{(i+1)} - \theta^{(i)}$

$\log w_\text{r}^{(i+1)} = \log p_\text{r}^{(i)}(\boldsymbol{y}') - \log p_\text{r}^{(i)}(\boldsymbol{y})$

$\log w_\text{o}^{(i+1)} = \log p_\text{o}^{(i+1)}(x_c) - \log p_\text{o}^{(i)}(x_c)$

$\log w^{(i+1)} = (1 - m_c)\log w_\text{r}^{(i+1)} + m_c \log w_\text{o}^{(i+1)}$ ▷ Apply mask (1/0 if $x_c$ known/unknown)

**return** $\log w^{(i+1)}$

**Algorithm 26** Particle logit continuous

**def** particleLogit( $\theta^{(i)}{=}\{\mu^{(i)}, \rho^{(i)}\}$, $\theta^{(i+1)}{=}\{\mu^{(i+1)}, \rho^{(i+1)}\}$, $\hat{\theta}^{(i)}{=}\{\hat{\mu}^{(i)}, \rho^{(i)}\}$, $\hat{\theta}^{(i+1)}{=}\{\hat{\mu}^{(i+1)}, \rho^{(i+1)}\}$, $y$, $\alpha$, $x_c$, $m_c$ ):

$p_\text{r}^{(i)} \leftarrow \mathcal{N}\left(\hat{\mu}^{(i)}, \alpha^{-1}\right)$ ▷ Receiver distribution

$p_\text{o}^{(i)},\, p_\text{o}^{(i+1)} = \mathcal{N}(\hat{\mu}^{(i)}, 1/\rho^{(i)}),\, \mathcal{N}(\hat{\mu}^{(i+1)}, 1/\rho^{(i+1)})$ ▷ Output distribution(s)

$y' = \frac{1}{\alpha}\left(\mu^{(i+1)}\rho^{(i+1)} - \mu^{(i)}\rho^{(i)}\right)$

$\log w_\text{r}^{(i+1)} = \log p_\text{r}^{(i)}(\boldsymbol{y}') - \log p_\text{r}^{(i)}(\boldsymbol{y})$

$\log w_\text{o}^{(i+1)} = \log p_\text{o}^{(i+1)}(x_c) - \log p_\text{o}^{(i)}(x_c)$

$\log w^{(i+1)} = (1 - m_c)\log w_\text{r}^{(i+1)} + m_c \log w_\text{o}^{(i+1)}$ ▷ Apply mask (1/0 if $x_c$ known/unknown)

**return** $\log w^{(i+1)}$

#### D. Auxiliary Algorithms

**Dynamic Time Warping** To obtain the optimal alignment between two sequences of points, $\mathbf{A} = \{(x_1, y_2, z_3), \ldots, (x_N, y_N, z_N)\}$ and $\mathbf{B} = \{(x_1, y_2, z_3), \ldots, (x_M, y_M, z_M)\}$, the pairwise distance/cost matrix $\mathbf{C} \in \mathbb{R}^{N\times M}$ is first calculated, where $\mathbf{C}_{i,j}$ is the distance between $\mathbf{a}_i \in \mathbf{A}$ and $\mathbf{b}_j \in \mathbf{B}$. The optimum warping path through $\mathbf{C}$, $\mathbf{P}$, is then computed using Algorithm 27. The length of $\mathbf{P}$, $L$, is then used to calculate the overall distance given the optimal alignment $d$ between $\mathbf{A}$ and $\mathbf{B}$, such that $d = 1/L\sum_{i=1}^{L}\mathbf{C}_{q_i}$ where $q_i \in \mathbf{P}$.

**Algorithm 27** Dynamic Time Warping

**def** dynamicTimeWarping($\mathbf{C} \in \mathbb{R}^{N\times M}$): ▷ Input: pairwise per-atom distance matrix, $\mathbf{C}$
$\mathbf{C}^A \leftarrow \mathbb{R}^{N\times M}$ ▷ Initialise empty accumulated cost matrix, $\mathbf{C}^A$
**while** $n \leq N$ **do** ▷ Fill first column of $\mathbf{C}^A$
    $\mathbf{C}^A_{n,1} \leftarrow \sum_{k=1}^{n} \mathbf{C}_{k,1}$
**while** $m \leq M$ **do** ▷ Fill first row of $\mathbf{C}^A$
    $\mathbf{C}^A_{1,m} \leftarrow \sum_{k=1}^{m} \mathbf{C}_{1,k}$
**for** $n = 2, ..., N$ **do** ▷ Fill the rest of $\mathbf{C}^A$
    **for** $m = 2, .., $ **do**
        $\mathbf{C}^A_{n,m} \leftarrow \mathbf{C}_{n,m} + \min\{\mathbf{C}^A_{n-1,m-1}, \mathbf{C}^A_{n-1,m}, \mathbf{C}^A_{n,m-1}\}$
$l \leftarrow 1, q_l \leftarrow (N, M)$
**repeat** ▷ Find the optimal path through $\mathbf{C}^A$
    $l \leftarrow l + 1, q_{l-1} \leftarrow (n, m)$
    **if** $n = 1$ **then**
        $q_l \leftarrow (1, m-1)$
    **else if** $m = 1$ **then**
        $q_l \leftarrow (n-1, 1)$
    **else**
        $q_l \leftarrow \text{argmin}\{\mathbf{C}^A_{n-1,m-1}, \mathbf{C}^A_{n-1,m}, \mathbf{C}^A_{n,m-1}\}$
**until** $q_l = (1, 1)$
$L \leftarrow l, \mathbf{P} \leftarrow \{q_L, q_{L-1}, ..., q_1\}$ ▷ Define optimal path, $\mathbf{P}$, and its length, $L$
**return** $L, \mathbf{P}, \mathbf{C}^A$

#### E. Supplementary Figures

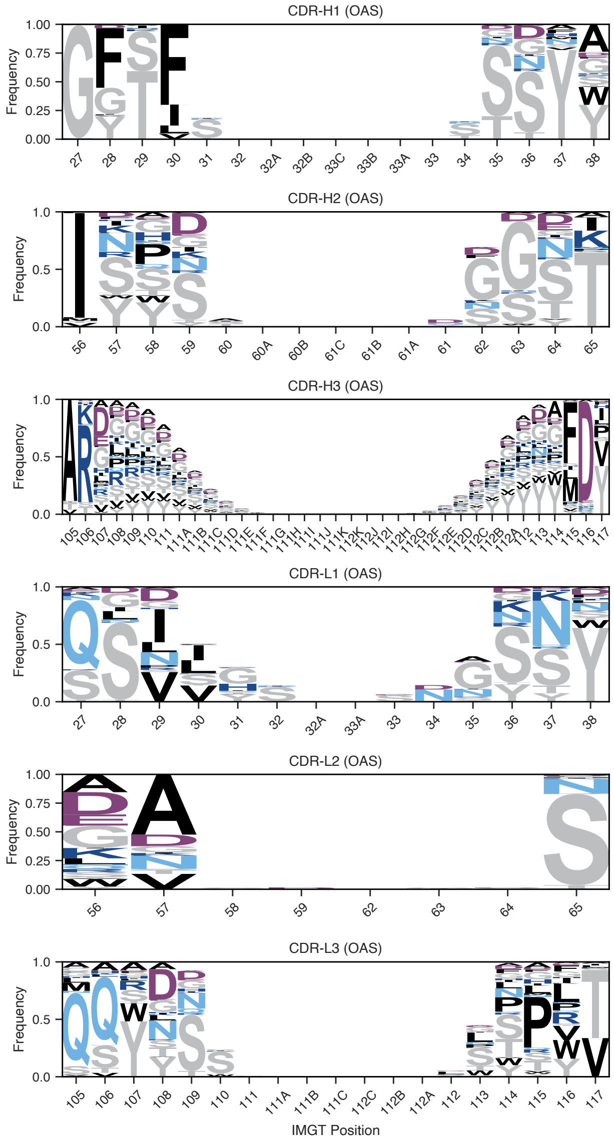

**Figure S1 | Position-specific amino acid propensities across the hold-out set.** Position-specific probability matrices were generated for the hold-out set by calculating normalised counts for each residue at each position. Gaps in the IMGT-aligned numbering were treated as a residue type. Residues are coloured according to their physicochemical properties.

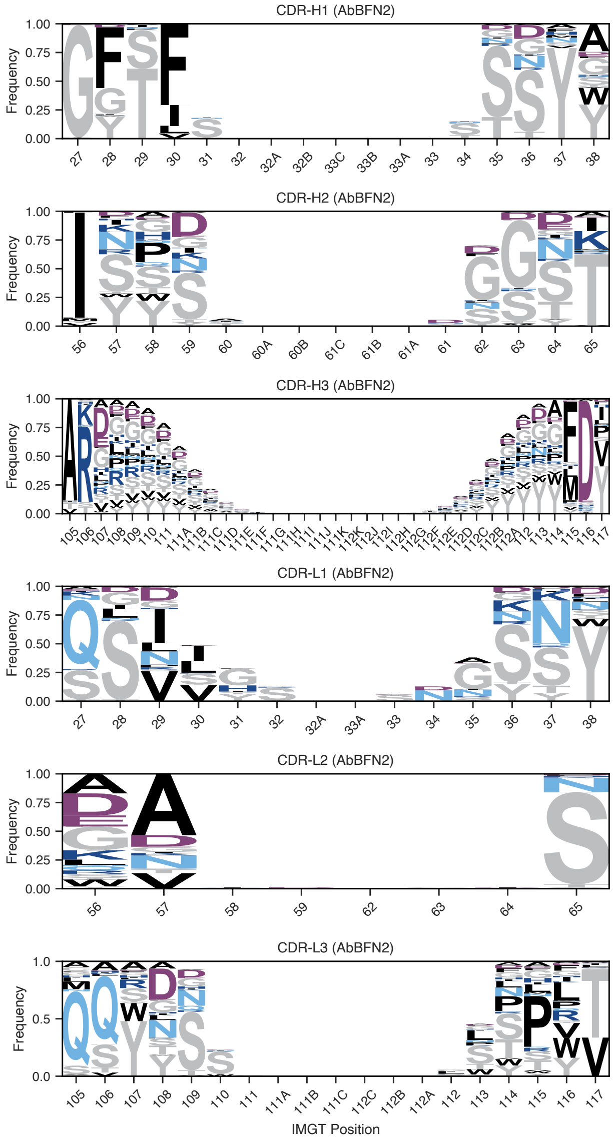

**Figure S2 | Position-specific amino acid propensities across generated samples.** Position-specific probability matrices were generated for 10,000 samples generated using AbBFN2 by calculating normalised counts for each residue at each position. Gaps in the IMGT-aligned numbering were treated as a residue type. Residues are coloured according to their physicochemical properties.

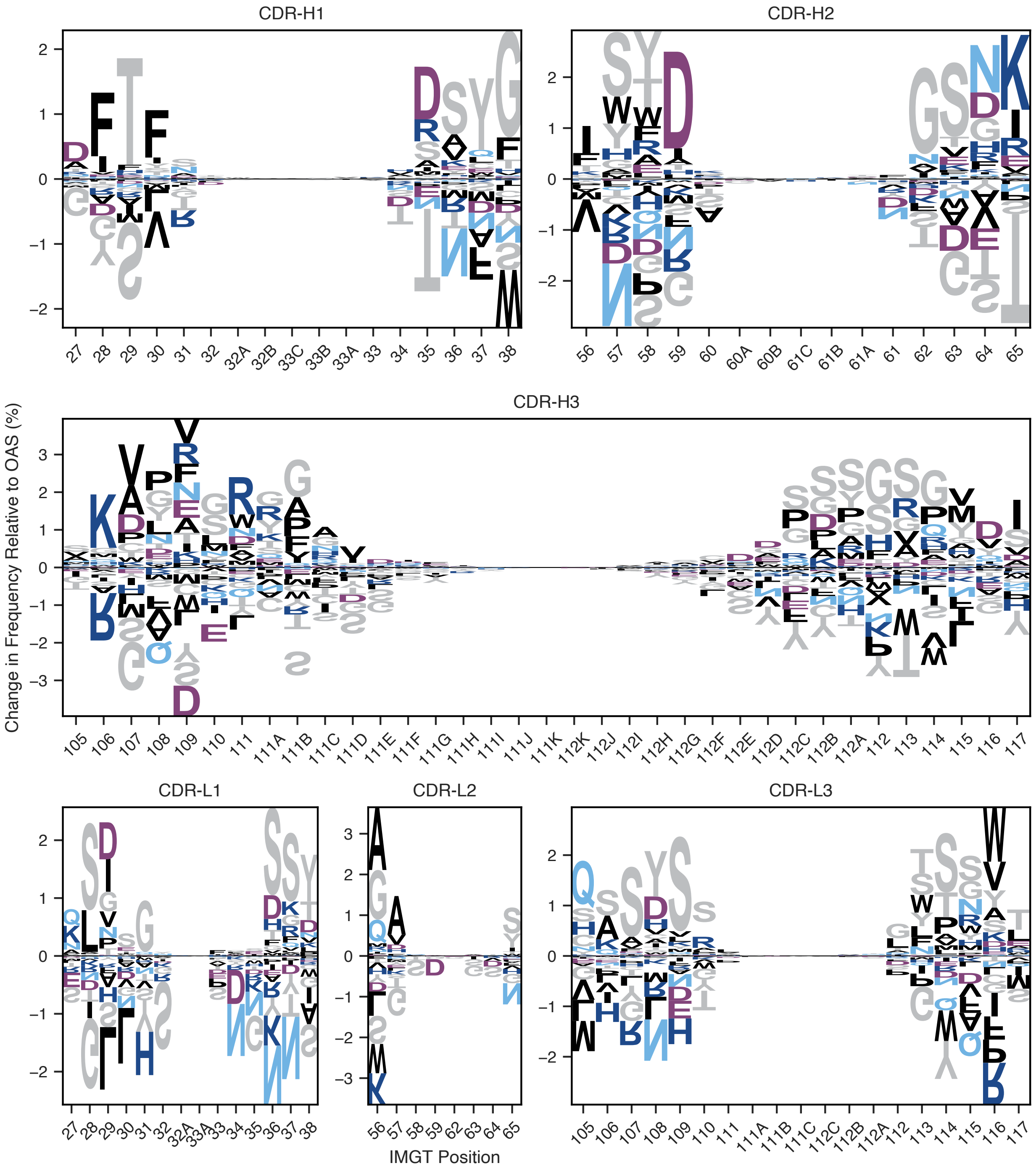

**Figure S3 | Logo plots of position specific residue propensity differences.** Position-specific deviations of amino acid frequencies of generated sequences when compared to the held-out data across CDR loops were calculating by taking the difference between position-specific probability matrices of the generated and held-out samples for each CDR loop. Residues are coloured according to their physicochemical properties. Positive values correspond to amino acids that are overrepresented in generated samples, and *vice versa*.

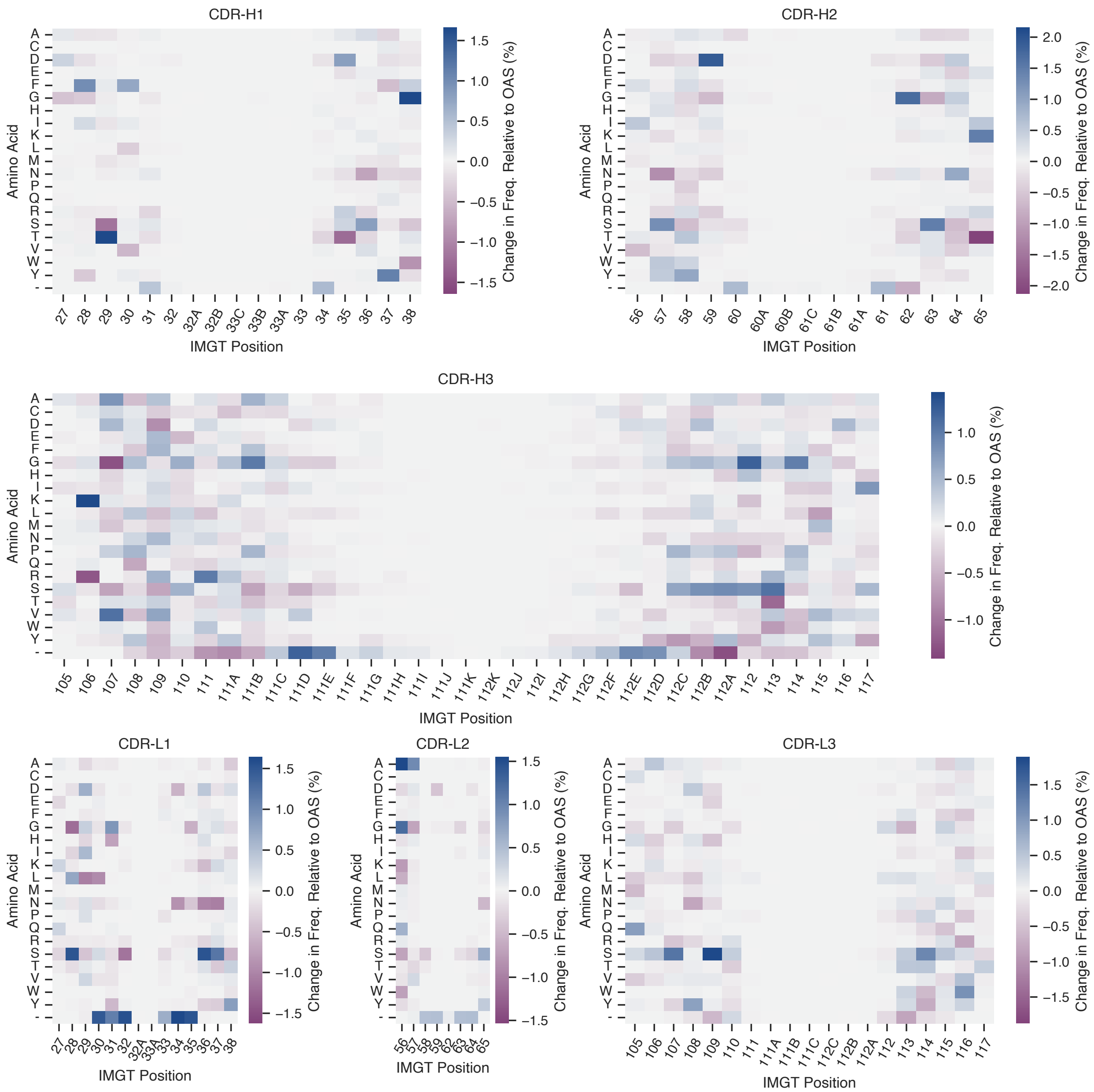

**Figure S4 | Heatmaps of position specific residue propensity differences.** Position-specific deviations of amino acid frequencies of generated sequences when compared to the held out data across CDR loops were calculating by taking the difference between position-specific probability matrices of the generated and held out samples for each CDR loop. Positive values correspond to amino acids that are overrepresented in generated samples, and *vice versa*.

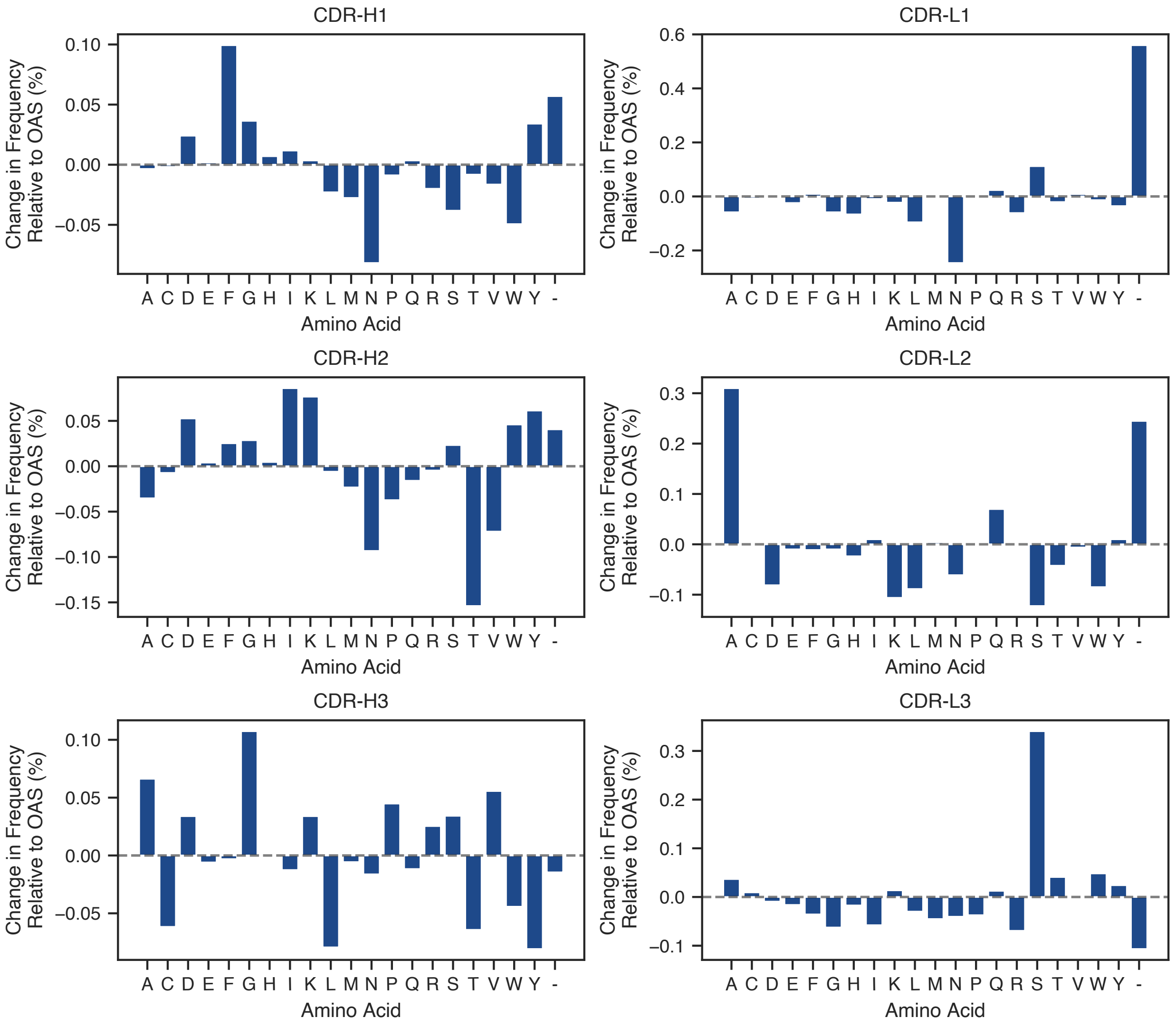

**Figure S5** | **Average differences in amino acid frequencies across loops.** For each loop (defined using the IMGT scheme), we calculated the average frequency of different amino acids across the entire loop for both held out and generated samples and calculated the differences. Positive values correspond to amino acids that are overrepresented in generated samples, and *vice versa*. "-" is used to represent gaps in the IMGT-aligned positions.

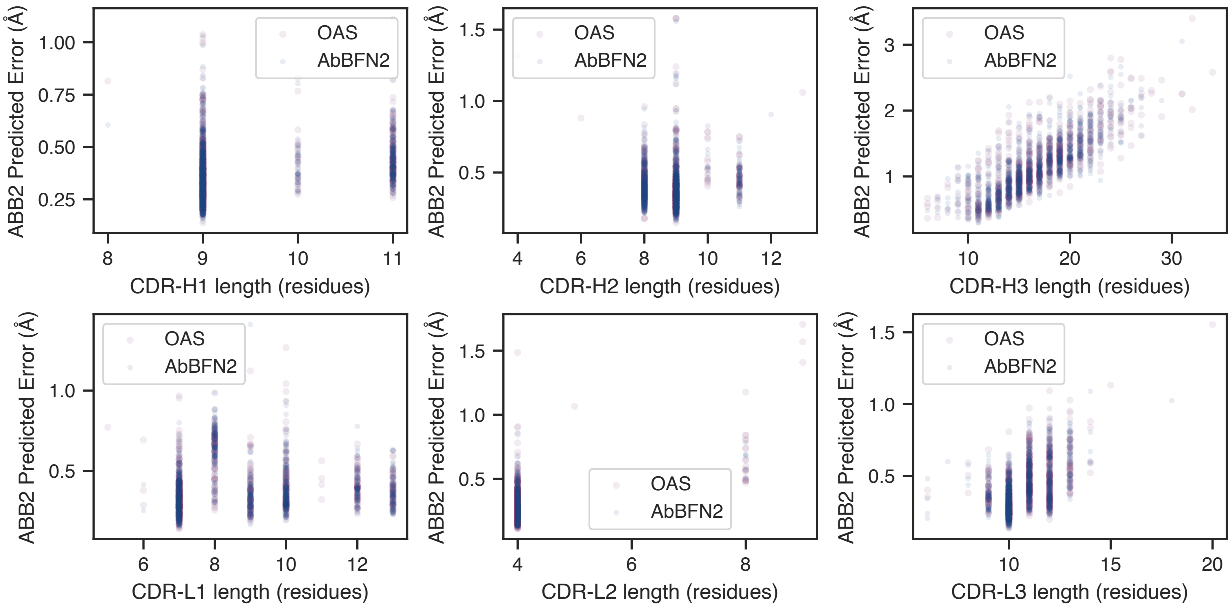

**Figure S6 | Relationships between sequence length and ImmuneBuilder predicted error scores.** For 9995 successfully modelled generated sequences (navy), and 10,000 held-out sequences (purple) and for each CDR loop, we calculated the average root mean squared predicted error across the entire loop. Results are shown for a random subset of 1,000 sequences from each set in order to avoid overcrowding the shown plots.

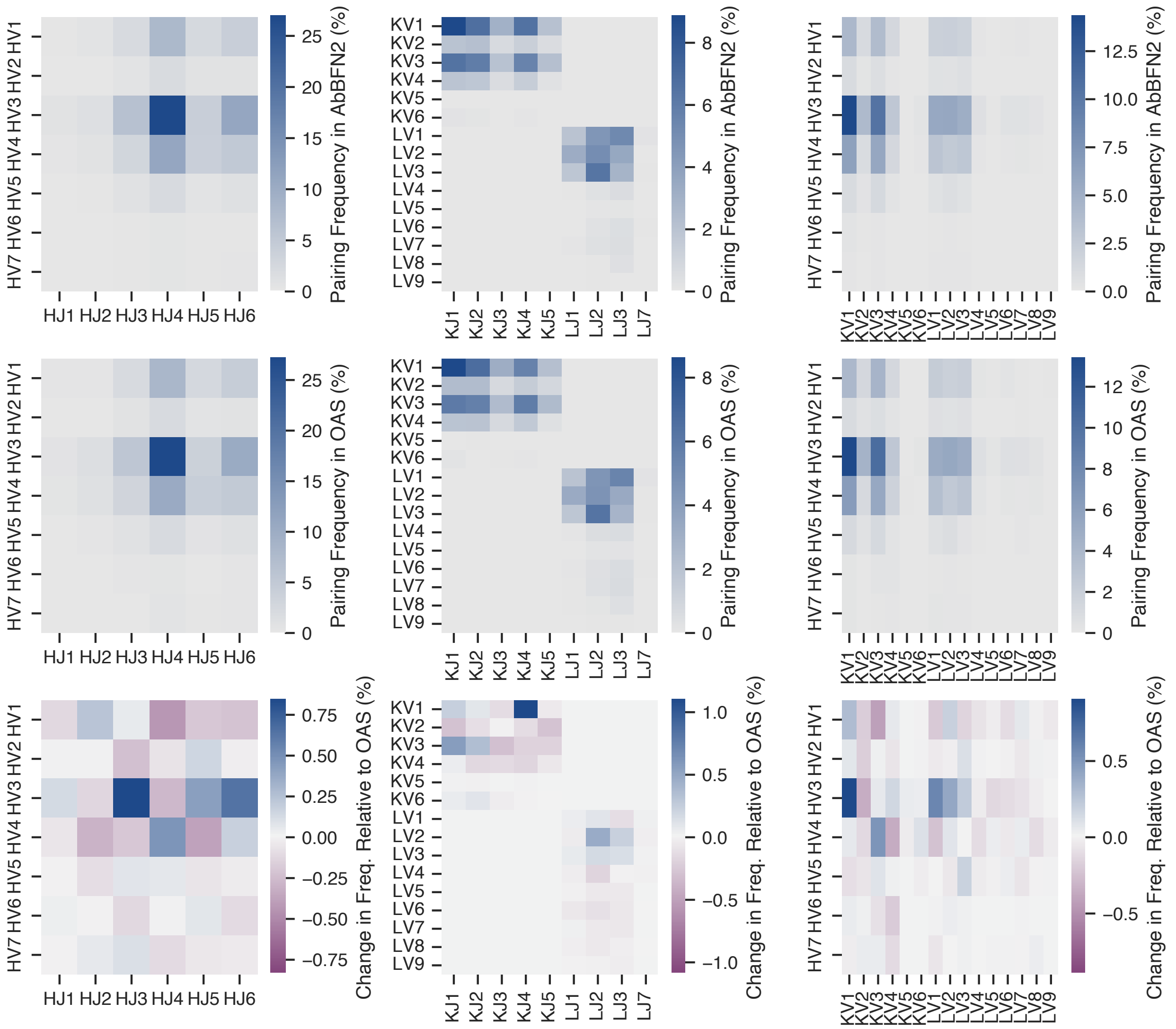

**Figure S7 | Frequencies of gene pairings and differences across generated and hold-out samples.** The top two rows show pairings across *V*- and *J*-genes of held out and generated samples. For the OAS-derived samples, the reported genes are the labels found in OAS. For generated samples. the reported labels are those output by AbBFN2. The bottom row shows the difference in frequencies across held-out and generated samples. Positive values correspond to amino acids that are overrepresented in generated samples, and *vice versa*.

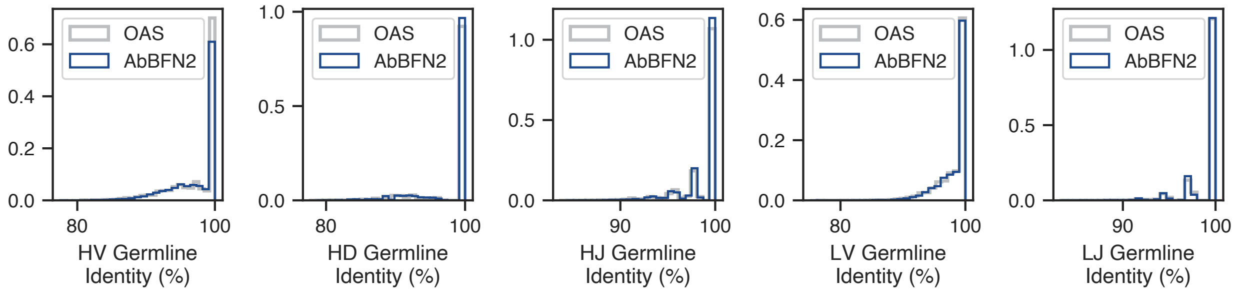

**Figure S8 | Distributions of sequence identities to germline genes.** The hold-out set identities are those derived directly from OAS. For AbBFN2-generated samples, we report the gene identity values generated by the model.

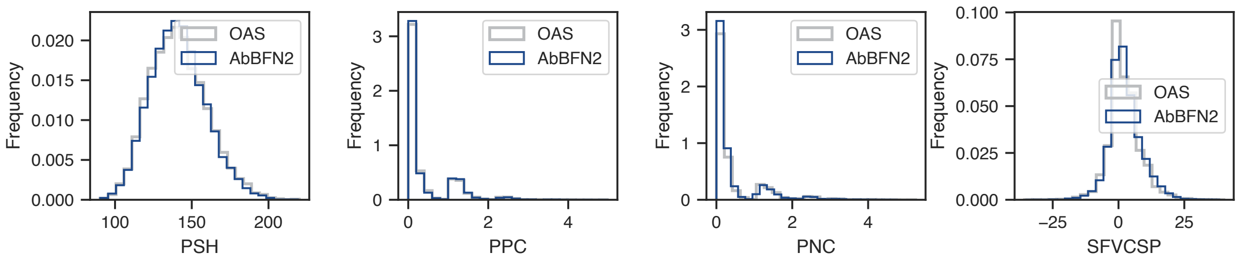

**Figure S9 | Distributions of Therapeutic Antibody Profiler (TAP) metrics.** TAP metrics were calculated for the OAS-derived hold-out set (folded using ImmuneBuilder) using an in-house implementation of TAP. For AbBFN2, the model-generated score is reported.

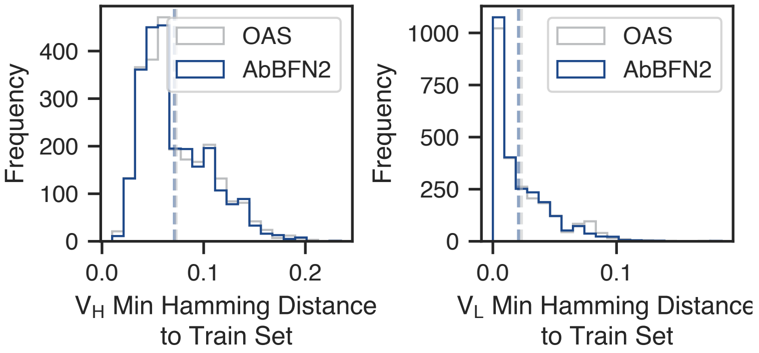

**Figure S10 | Minimum normalised Hamming distances of the held out and generated samples to the training set.** Hamming distances between the held-out data (grey) and generated samples (navy) and a random sample of 10,000 sequences from the training set. Briefly, each sequence was aligned and padded to the most commonly found 200 IMGT positions, before distance calculation and normalisation. For each generated or held-out sequence, the smallest Hamming distance to the train set is reported. Dashed lines correspond to means across all sequences. The differences between means is significant only for $V_L$ sequences ((Mann-Whitney $U = 3014736.0, p = 0.03$))

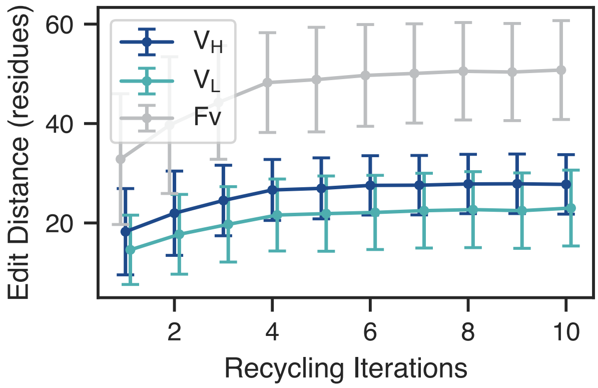

**Figure S11 | Number of total mutations accumulated during sequence humanisation.** Starting with 25 precursor sequences of experimentally humanised clinical stage therapeutics, we performed sequence humanisation with a total budget of 10 recycling iterations per sequence. After each iteration, we calculated the edit distance between the current design and starting sequence. The lineplots show means across all samples, with error bars showing standard deviations.

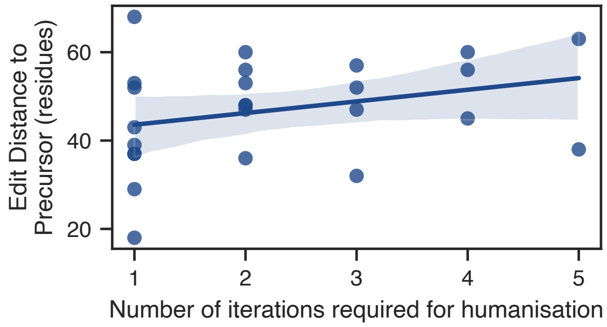

**Figure S12** | **Number of mutations and steps required for sequence humanisation.** Starting with 25 precursor sequences of experimentally humanised clinical-stage therapeutics, we performed sequence humanisation with a total budget of 10 recycling iterations per sequence. For each sequence, we then extracted the first instance in the recycling procedure where the probability of the sequence being human under the model was greater than or equal to 0.95 and calculated its edit distance to the precursor sequence. The line of best fit is shown, with the shaded area corresponding to the 95% confidence interval, calculated using bootstrapping.

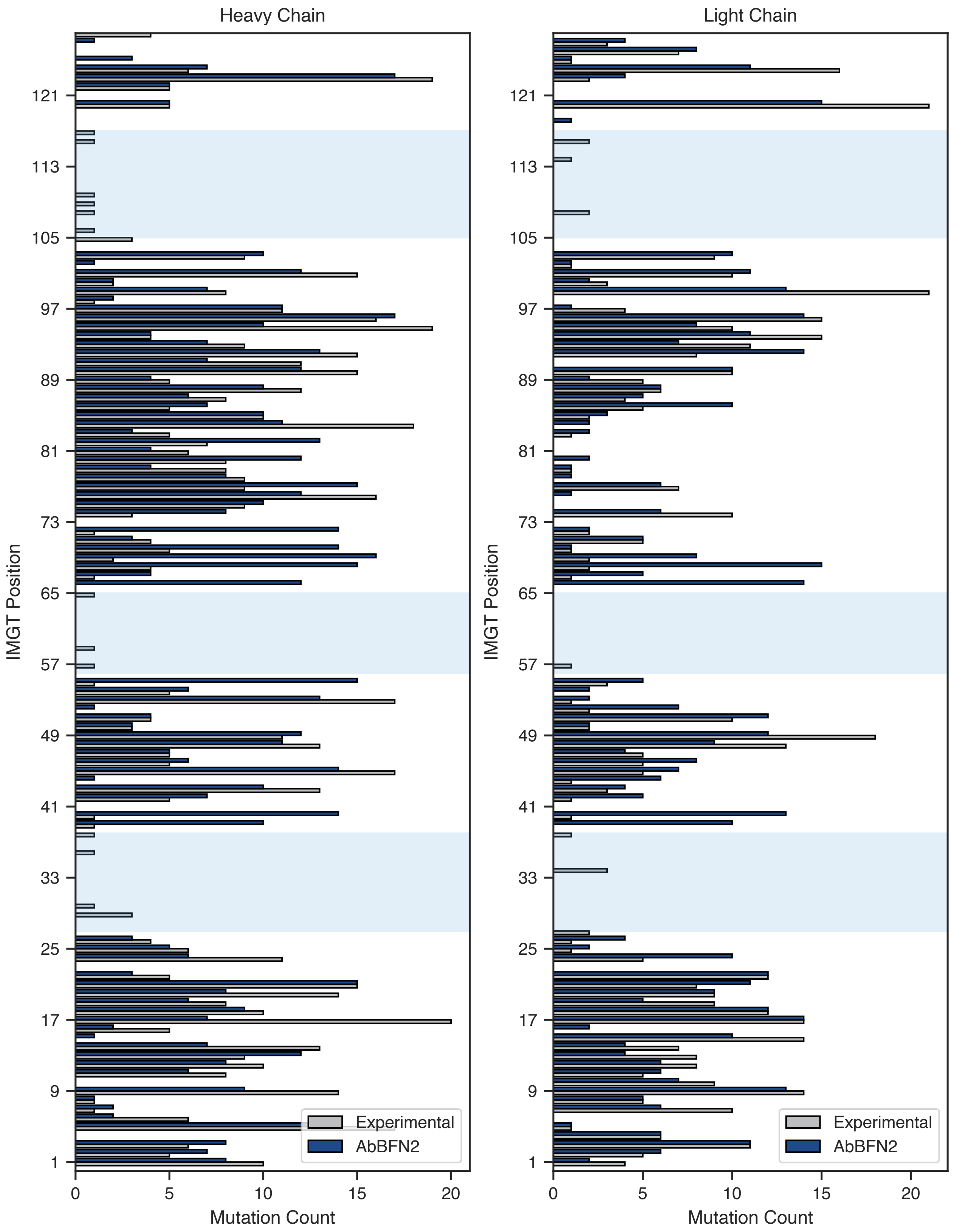

**Figure S13** | **Position-specific mutations in experimentally and computationally humanised sequences.** Starting with 25 precursor sequences of experimentally humanised clinical stage therapeutics, we performed sequence humanisation with a total budget of 10 recycling iterations per sequence. Here, we show the number of times an IMGT position has been mutated in both our samples and the experimentally humanised sequences. Light blue shaded areas correspond to CDR loops.

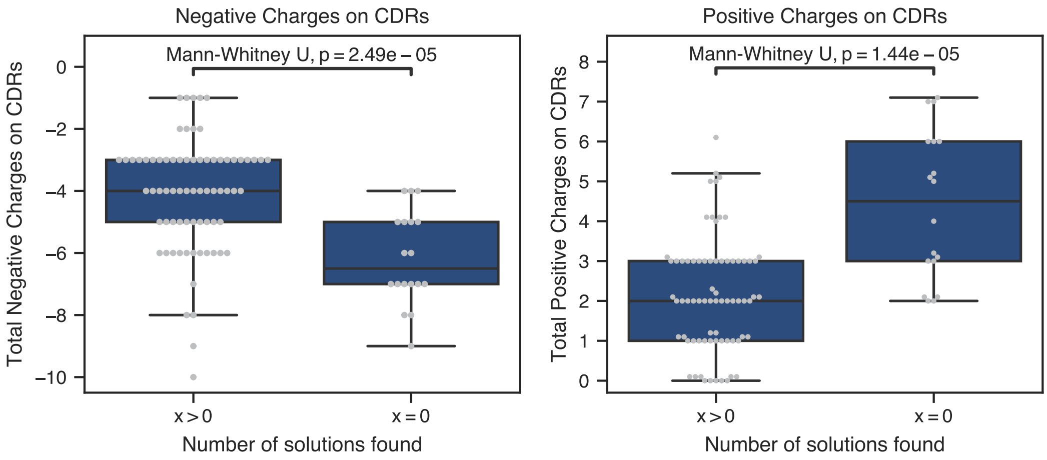

**Figure S14 | Charges of CDR residues in successful and unsuccessful optimisation trajectories.** Of the 91 non-human sequences with development liabilities that were optimised, $n = 73$ had at least one valid solution that AbBFN2 identified ($x > 0$), while for $n = 18$, AbBFN2 did not generate valid solutions ($x = 0$). We calculated the total charge of CDR residues in these sequences using the TAP definitions of charged residues (Asp, Glu: -1; Arg, Lys: +1; His: +0.1) and performed Mann-Whitney U tests for these. Boxplots show the 25%, 50% (median), and 75% quartiles. Whiskers extend to a maximum of 150% of the interquartile range.

### F. Supplementary Results

#### F.1. Extended analysis of unconditionally generated CDR loops

As the CDR loops are the most functionally relevant and variable regions of an antibody, we examined the CDR lengths of the generated sequences (Fig. 2a-e) and found excellent agreement to natural sequences, with only very small deviations. Notably, the AbBFN2-generated sequences faithfully capture the distribution of lengths of the CDR-H3 loop (Fig. 2c) which is the most variable CDR loop and often the major mode of antigen recognition. We also examined the individual sequences of the CDR loops by calculating residue propensities across IMGT positions [8] for each loop (Fig. S1, Fig. S2), as the IMGT numbering scheme aims to assign numbering based on functional similarity. In order to find specific positions where a residue is under- or overrepresented in our samples, we also calculated the difference between the position probability matrices (PPMs) for generated and held out samples (Fig. S3, Fig. S4, Fig. 2g). We found that deviations were no greater than $1.5\,\%$, and were often a result of amino acids being replaced by similar residues. For example, IMGT position 29 in generated $V_H$ sequences tend to have a roughly $1.5\,\%$ enrichment in Thr residues compared to Ser residues, both of which are small and hydrophilic (Fig. S3). Similarly, $V_H$ position 106 (on the CDR-H3 loop) is slightly enriched in Lys residues in favour of Arg residues, both of which are positively charged (Fig. 2g, Fig. S3). Upon marginalising out the positions (i.e. examining whether a residue is under- or overrepresented across the entire CDR), we found that the deviations were even smaller, with a maximum magnitude of approximately $0.3\,\%$ (Ser residues in CDR-L3) (Fig. S5).

#### F.2. CDR Sequence Inpainting Using AbBFN2

We tested the ability of AbBFN2 to recover each CDR loop given the remainder of the antibody sequence (Table S3). As expected, we found that AbBFN2 has lower AAR rates when compared to two leading BERT-style protein language models, AntiBERTy [9] and AbLang2 [10]. We hypothesize that this is due to two factors, namely the limited training set when compared to these models, as well as the the data splitting procedure. The former factor contributes in the form of observed diversity. All other models were trained on the entirety of the OAS database, where paired sequences (2M) are vastly outnumbered by unpaired sequences (2.5B). Due to the sequence diversity exhibited in antibody repertoires, this means that AbBFN2 is exposed to far fewer sequences during training. Considering the train-test-validation split, we note that although we followed the same splitting scheme as AbLang2, the exact sequences which fall into each split are not identical, meaning that most of our held out sequences will have been part of the training set for all other models.

**Table S3 | Performance on sequence inpainting tasks.** For each CDR loop, the IMGT residues corresponding to that loop were masked out and predicted, with the remainder of the sequence provided as conditioning information. For AbBFN2 and AbLang2, both the heavy and light chain sequences were provided. For AntiBERTy, only the relevant chain was used. For AbBFN2, different sets of conditioning information are shown in brackets.

| Model | Context | CDR-H1 | CDR-H2 | CDR-H3 | CDR-L1 | CDR-L2 | CDR-L3 |
|---|---|---|---|---|---|---|---|
| AbLang2 | - | **91.23** | **89.57** | **45.35** | **91.46** | **91.64** | **85.84** |
| AntiBERTy | - | 90.35 | 88.07 | 43.23 | 90.02 | 89.95 | 83.82 |
| AbBFN2 | Seq. only | 85.69 | 82.89 | 34.46 | 86.62 | 87.21 | 79.04 |
| AbBFN2 | Seq. + $V_H$ *V*-gene | 85.88 | 82.83 | - | - | - | - |
| AbBFN2 | Seq. + $V_H$ *V,J*-genes | - | - | 36.94 | - | - | - |
| AbBFN2 | Seq. + $V_H$ *V,D,J*-genes | - | - | 38.15 | - | - | - |
| AbBFN2 | Seq. + $V_L$ *V*-gene | - | - | - | 86.76 | 87.3 | - |
| AbBFN2 | Seq. + $V_L$ *V,J*-genes | - | - | - | - | - | 80.01 |

##### F.3. Assessment of implicitly modelled properties of VRC-01-like antibodies

During conditional generation, it is important to ensure that relationships between variables that have not been conditioned on are appropriately shifted based on the conditioning information. For instance, when conditioning on a specific $V_H$ *V*-gene, the CDR-H1 and CDR-H2 sequences should reflect the underlying condiitonal distribution. To this end, we examined a number of dependencies that might be influenced by the conditioning information. We analysed the spread of germline sequence identities to the IGHV1-2 germline sequence using both predictions by the model, and ANARCI as validation (Fig. 5k) and find that the generated sequences do not simply recapitulate the requested germline sequence, but match the natural distribution of germline sequence identities (Fig. S8).

Next, we compared the $V_L$ *V*-gene family distribution of our VRC-01-like library to that of similar sequences (correct species, light chain locus, $V_H$ *V*-gene, and a CDR-L3 length of less than 9 [we included all short CDR-L3 loops, since there were not enough loops of exactly 5 residues]) in the hold-out set. Here, we find that the $V_L$ *V*-gene families are appropriately captured, with the majority of samples belonging to IGKV3/1 ($74\,\%$ for hold-out, $93\,\%$ for generated VRC-01-like samples), a smaller number coming from the IGKV4/2 lineages ($26\,\%$ for hold-out, $7\,\%$ for generated VRC-01-like samples), and a small remainder coming from the IGKV6 lineage.

We also assessed the diversity of the generated samples, and found 149 unique CDR-H1 sequences, 147 unique CDR-L2 sequences, and 1715 unique CDR-H3 sequences. This trend matches the expected diversity. CDR-H3 is both more diverse generally and has contributions from all three germline genes, whereas CDR-H1/H2 only have sequence contributions from the *V*-gene. Since this gene is one of the explicit conditions during sampling, we find less diversity amongst these loops, whereas free sampling for most of the CDR-H3 sequence is allowed in this experiment. Furthermore, we find a broad distribution of CDR-H3 loop lengths, with an average of $15.9 \pm 3.1$, matching the background distribution (Fig. 2c).
